# Behavioral sequences across multiple animal species in the wild share common structural features

**DOI:** 10.1101/2024.01.20.576411

**Authors:** Pranav Minasandra, Emily M Grout, Katrina Brock, Margaret C Crofoot, Vlad Demartsev, Andrew S Gersick, Ben T Hirsch, Kay E Holekamp, Lily Johnson-Ulrich, Amlan Nayak, Josué Ortega, Marie A Roch, Eli D Strauss, Ariana Strandburg-Peshkin

## Abstract

Animal behavior can be decomposed into a sequence of discrete activity bouts over time. Analyzing the statistical structure of such behavioral sequences can provide insights into the drivers of behavioral decisions. Laboratory studies, predominantly in invertebrates, have suggested that behavioral sequences exhibit multiple timescales and long-range memory, but whether these results can be generalized to other taxa and to animals in natural settings remains unclear. By analyzing accelerometer-inferred predictions of behavioral states in three species of social mammals (meerkats, white-nosed coatis, and spotted hyenas) in the wild, we discovered surprisingly consistent structuring of behavioral sequences across all behavioral states, all individuals, and all study species. Behavioral bouts were characterized by decreasing hazard functions, wherein the longer a behavioral bout had progressed, the less likely it was to end within the next instant. The predictability of an animal’s future behavioral state as a function of its present state always decreased as a truncated power-law for predictions made farther into the future, with very similar estimates for the power law exponent across all species. Finally, the distributions of bout durations were also heavy-tailed. Why such shared structural principles emerge remains unknown, and we explore multiple plausible explanations, including environmental non-stationarity, behavioral self-reinforcement, and the hierarchical nature of behavior. The existence of highly consistent patterns in behavioral sequences across our study species suggests that these phenomena could be widespread in nature, and points to the existence of fundamental properties of behavioral dynamics that could drive such convergent patterns.

**Significance statement:** The study of animal behavior seeks to understand how and why animals do what they do. This pursuit of general principles governing behavior across species can be approached by first understanding *when* animals choose to change their behavioral states (e.g., switching from walking to standing, or to running). Using accelerometer-inferred behaviors of three social mammals, we uncover common structural ‘long timescale’ patterns in their sequences of behavior. We explore two explanations, involving either positive feedbacks or the interaction of several independent time-scales, about how such common patterns arise.

## 1 Introduction

A day in the life of an animal consists of a series of transitions between discrete behavioral states – for example, a meerkat might wake up, stand in the sun outside its burrow as the air temperature rises, then eventually move off in search of food, occasionally scanning the skies for predators. When combined, these actions describe a behavioral sequence. The behavior of all organisms, from oomycetes to orcas, consists of such behavioral sequences, and these sequences in turn reflect complex processes in the animal brain that, together with environmental information, underlie decisions about which behavior to perform at any moment in time [1]. Characterizing the statistical properties of behavioral sequences and, in particular, quantifying the time-scales that structure them [e.g., 2, 3], can shed light on the underlying decision-making processes that generate observed patterns of behavior.

It is likely that some aspects of these decision-making processes are shared across species, either because of common evolutionary origins or convergent evolution. Searching for common patterns in the statistical structure of behavioral sequences is therefore a powerful tool for identifying potential general principles of animal behavior.

Several previous studies have pointed towards the existence of a large number of operant time-scales in the organization of behavior [2, 4–8], such that behavior acts as a process with long-range memory. Taken together, work on different species suggests the potential for general, widespread structural principles in behavioral sequences, but no studies have yet applied these analytical approaches to empirical data on multiple types of behavior in multiple species in natural settings to directly assess the shared structural features (if any) of behavioral sequences.

Here, we use continuous accelerometer data recorded via tracking collars to infer behavioral sequences across three species of social mammals – 15 meerkats (*Suricata suricatta*; small burrow-living mongooses), 9 white-nosed coatis (*Nasua narica*; fruit-eating omnivores), and 5 spotted hyenas (*Crocuta crocuta*; large carnivores). Applying three analytical approaches to analyze data from each species, we find shared structural properties across all species, behaviors, and individuals, indicating common features in the underlying decision-making processes at play. We hypothesize that such common decision-making processes might be the result of equally common drivers of behavior: non-stationarity due to animals responding to constantly-varying external variables, or a positive-feedback-centered decision-making structure as a feature of behavioral algorithms.

### 1.1 Characterizing behavioral sequences

There are many ways to characterize a behavioral sequence. Here we focus on three approaches: measuring hazard functions, characterizing bout duration distributions, and quantifying the decay of mutual information between present and future behavioral states. These three methods provide complementary perspectives on the dynamical structure of behavior.

#### Behavioral hazard functions

A bout is a process that has a defined start and end time. In the case of behavior, a bout defines a contiguous period of time during which an animal performs the same behavior, for example walking, resting, or foraging. Survival analytic concepts [9, 10] are useful in understanding the nature of processes with observable beginnings and ends. Of these concepts, specifically useful is the hazard function, *h*(*t*), which quantifies the instantaneous probability of a process (in this case a behavior) ending given that it has already been in progress for time *t*. I.e., if *τ* is the survival time of a behavioral bout from start to end, then

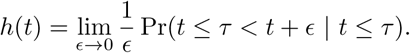

Qualitatively, an increasing hazard function represents a process that is wearing out, becoming more likely to end as time passes [e.g., the survival of leukemia patients, 11]. Conversely, a decreasing hazard function represents a process that is reinforced, becoming progressively less likely to end as time passes, such as the survival of patients recovering from surgeries [11] or the operational durations of spacecraft [12]. Constant (i.e., flat) hazard rates are associated with memory-less processes, such as the radioactive decay of an unstable atomic nucleus. Many commonly used models of behavior and movement, such as simple Markov models, assume memory-less processes, where individuals switch between behaviors with constant transition probabilities dependent only on their current state. Such Markovian assumptions are often made for the sake of analytical tractability despite the fact that animal behavior is often non-Markovian [13].

The shapes of hazard functions can give information about the underlying behavioral processes. For instance, if the hazard function for a particular behavior was found to decrease up to some time *T* and then increase after that, we might predict that this timescale *T* is important to the animal’s decision-making process. The shape of the hazard function can provide insights into the nature of memory in the behavioral sequence—for instance, decreasing hazard functions might link to self-reinforcing processes that have strong long-range order [14].

*A priori*, we expect different behaviors to show bouts characterized by different types of hazard functions. For instance, behaviors that are energetically expensive or risky might have bouts with increasing hazard rates, since the animal becomes progressively less likely to continue such behaviors the longer they have persisted. Conversely, behaviors with increasing returns, such as extractive foraging from a complex food resource (such as a carcass or fruit with a tough exterior [15], or a termite mound [16]) that progressively yields higher quality nutrition as the animal forages, might be expected to have decreasing hazard rates up to the point where returns start to diminish. Since situations of diminishing marginal returns are likely to be more common in nature than those with increasing returns, we might expect that hazard functions should most often increase as the individual engages in the behavior to accomplish a certain task or meet a certain need; and therefore becomes more likely to switch to the next state as time progresses. This seems to be a common expectation. For instance, Gygax et al [10] write about sleep bouts: “this likeliness [i.e., the hazard function] may increase the longer the animal has been in the state”.

#### Mutual information decay

Information theoretic perspectives can reveal the timescales over which behavioral sequences show memory [17]. Predictivity measures the extent to which knowledge of the animal’s current behavioral state is predictive of its future state (conversely, the extent to which an animal’s past behavior influences its current state). The predictive value of the current state decays as one moves forward in time (i.e., it is increasingly difficult to forecast behavior further into the future).

Quantifying the rate at which the future’s dependence on the present state decays in an animal’s behavioral sequence can be informative about underlying decision-making processes [18, 19]. If predictivity decays exponentially, we could identify a characteristic timescale in the behavioral sequence at which behavioral algorithms operate. On the other hand, a ‘scale invariant’ power-law decay would indicate the interaction of processes acting at (potentially infinitely) many timescales [17, 20] that together generate behavior such that an animal’s present state is informative very far into the future. For instance, experiments on *Drosophila* have revealed long-term order in the fruit-flies’ behavioral sequences, interpreted as the result of behavioral decision-making processes acting at multiple time scales [2, 13, 21].

#### Behavioral bout duration distributions

Hazard functions of processes are linked directly to the distributions of process survival times (here, the durations of bouts of each behavior), such that knowing one provides information about the other. The durations of behavioral bouts are sometimes heavy-tailed (see references below). ‘Heavy-tail’ refers to a tail (the part of the distribution associated with large values, e.g., long bouts) which decays slower than an exponential distribution. Heavy-tailed distributions are characterized by an unusually high occurrence of very large values. Some examples of heavy-tailed distributions are the log-normal, stretched exponential, power-law, and truncated power-law distributions. If a variable is normally distributed, then a typical value, such as the mean, can convey useful information about the distribution of this variable. For example, if the time an animal spends in a given bout of vigilance were normally distributed with a mean of 10 seconds, it would imply that the timescale of 10 seconds is behaviorally relevant for that animal – perhaps a typical scan of the environment takes 10 seconds to complete. Because of the very high variances of heavy-tailed distributions, they are difficult to characterize with such single ‘typical’ values. In the case of behavioral bouts, a heavy-tailed distribution would imply that it is difficult to choose a single time scale at which behavioral decisions can be presumed to occur.

Such scale invariance is associated with power-law bout duration distributions (probability density function *f_τ_* (*t*) ∝ *t^−α^*). Penguin [22] and macaque [23] foraging patterns, *Drosophila* flight sequences [24] and activity patterns [25], as well as mouse resting behavior [26, 27] show scale invariance. Moreover, power-law distributed step lengths have been reported in movement trajectories of numerous species [the Lévy Flight Foraging Hypothesis, 28, 29], which indicate a power-law bout duration distribution for movement states. Truncated power-laws show power-law properties up to some cutoff, beyond which they behave similarly to exponential distributions (*f_τ_* (*t*) ∝ *t^−α^e^−λt^*). These distributions are often associated with phenomena with memory and positive feedback (e.g. self-reinforcement): the distributions of wait-times between certain events is often power-law distributed: e.g., the duration between earthquakes [30], neuronal avalanches [31], armed human conflict [32], or between consecutive e-mails [33]. Bouts of inactive behaviors (e.g., waiting for an event to occur) are theorized to be power-law distrubuted [33], and power-law distributions are empirically found in resting human patients [27, 34, 35] and other species such as mice [27]. Elsewhere, behavioral bouts have also been found to be lognormally distributed [36–38]. Lognormal distributions are also heavy-tailed, easily generated by the multiplication of positive random variables [39, 40], and have decreasing hazard functions at some parameter values.

## 2 Results

Deploying accelerometers on individuals of three species, we used either conventional or machine-learning based methods to quantify individuals’ behavioral sequences. Based on feasibility, behavioral sequences were generated at varying levels of precision while distinguishing different behaviors— coati behavioral sequences were composed of high vs low activity, whereas hyena and meerkat behaviors were more precisely identifiable, so that their sequences were composed of categories of kinematic states such as standing, walking, vigilance, and foraging (see Methods).

### 2.1 Decreasing hazard functions in bouts of all behaviors

From each individual’s sequence, we extracted the durations of all bouts of each behavioral state. From these durations, we quantified the hazard function for bouts of each behavior.

In contrast to our *a priori* expectations, we found that hazard functions were always decreasing, across all behavioral states, all individuals, and all species (Figure 1). This result implies that the longer an animal has been in a given behavioral state, the less likely it is to switch to a new state within the next instant.

**Figure 1:**
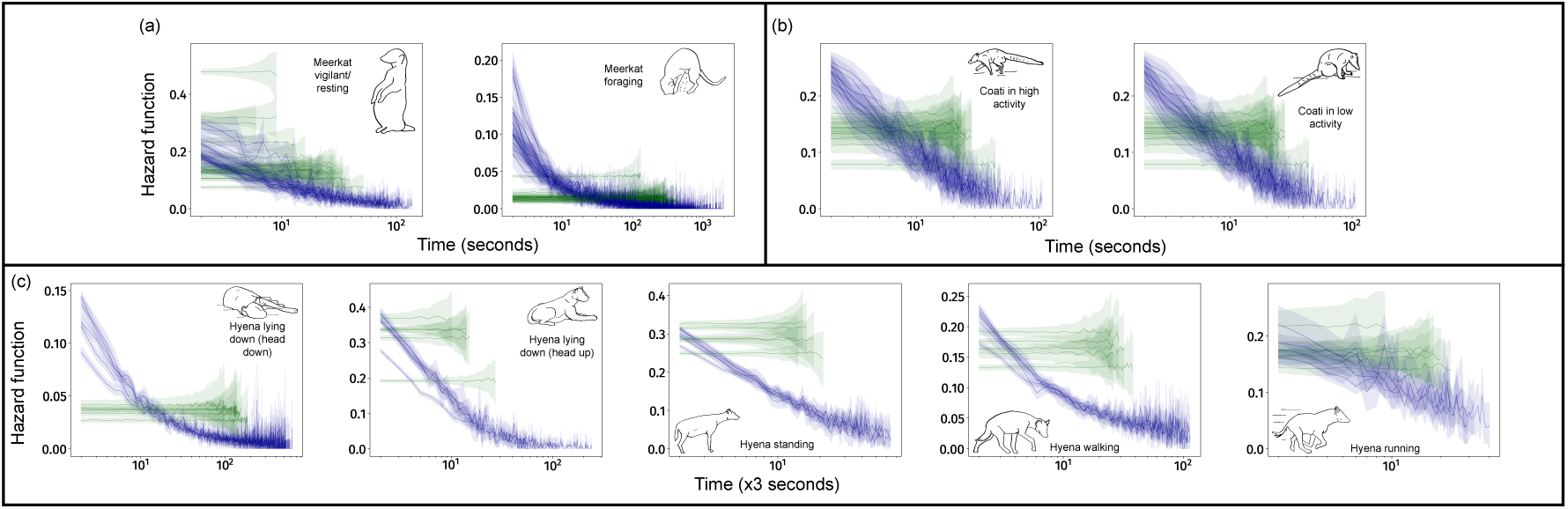
Decreasing hazard functions (blue lines) characterize the bouts of all behavioral states in (a) meerkats, (b) coatis, and (c) hyenas. Each blue line indicates the hazard function of bouts of a single individual, and shaded regions represent 95% Confidence Intervals. Green lines are average hazard functions and average 95% confidence intervals for thirty Markovian pseudosequences generated from the true data for each individual. These lines preserve instant-to-instant dynamics, and destroy longer time-scale dynamics in behavior. As expected from memorylessness, these hazard functions are flat in all cases. All hazard functions are visualized only up to the length of the 100th-longest bout. For each plot, inset drawings and label indicate the behavior in question. Hazard functions are decreasing across all species, all individuals, and all behaviors. Bouts of the meerkats’ running behavior are rare and short, making the hazard function impossible to quantify, and thus this state is not included. *x*-axes indicate time (in number of time-windows) spent in a bout of a given behavior. Behavioral sequences are inferred at resolutions of 1 s for meerkats and coatis, and 3 s for hyenas.

We also generated 30 pseudosequences based on the true behavioral sequences of each individual, so that the pseudosequences preserved only instant-to-instant (i.e., Markovian) aspects of behavioral dynamics, but eliminated dynamics at longer timescales. As expected in such purposefully memoryless data, hazard functions of all behaviors in all these pseudosequences were flat.

### 2.2 Long term predictivity of behavioral sequences

To identify characteristic timescales of behavioral algorithms, we measured predictivity decay in the behavioral sequences by asking how long into the future the current behavior of an animal remains predictive. To quantify this *predictivity*, for each individual, we computed the adjusted mutual information [41] between its behavioral state at time *t* and time *t* + *τ*, accounting for finite-size corrections [42].

In all three species, we found that the mutual information between behavioral states decays over time *τ* as an exponentially truncated power-law (Figure 2), with a surprisingly conserved power-law exponent (meerkats: 0.195 ± 0.06, coatis: 0.158 ± 0.015, hyenas: 0.187 ± 0.014; all variation here is standard deviation measured across individuals, SI Table S3 A-C). Interestingly, these exponents are also strikingly similar to the scaling dimension found recently in *Drosophila* behavioral sequences [21, 0.180 ± 0.005]. The timescale of truncation (i.e. the transition from power law to exponential decay) was consistent across all individuals within a species: at ∼ 1000 s (or ∼ 15 minutes) in all meerkats and coatis and ∼ 3000 s (∼ 45 min) in all hyenas. Moreover, the predictivity decay was far slower than in case of the Markov pseudosequences generated based on true data, showing the long time-scale dynamics of behavior in these animals.

**Figure 2:**
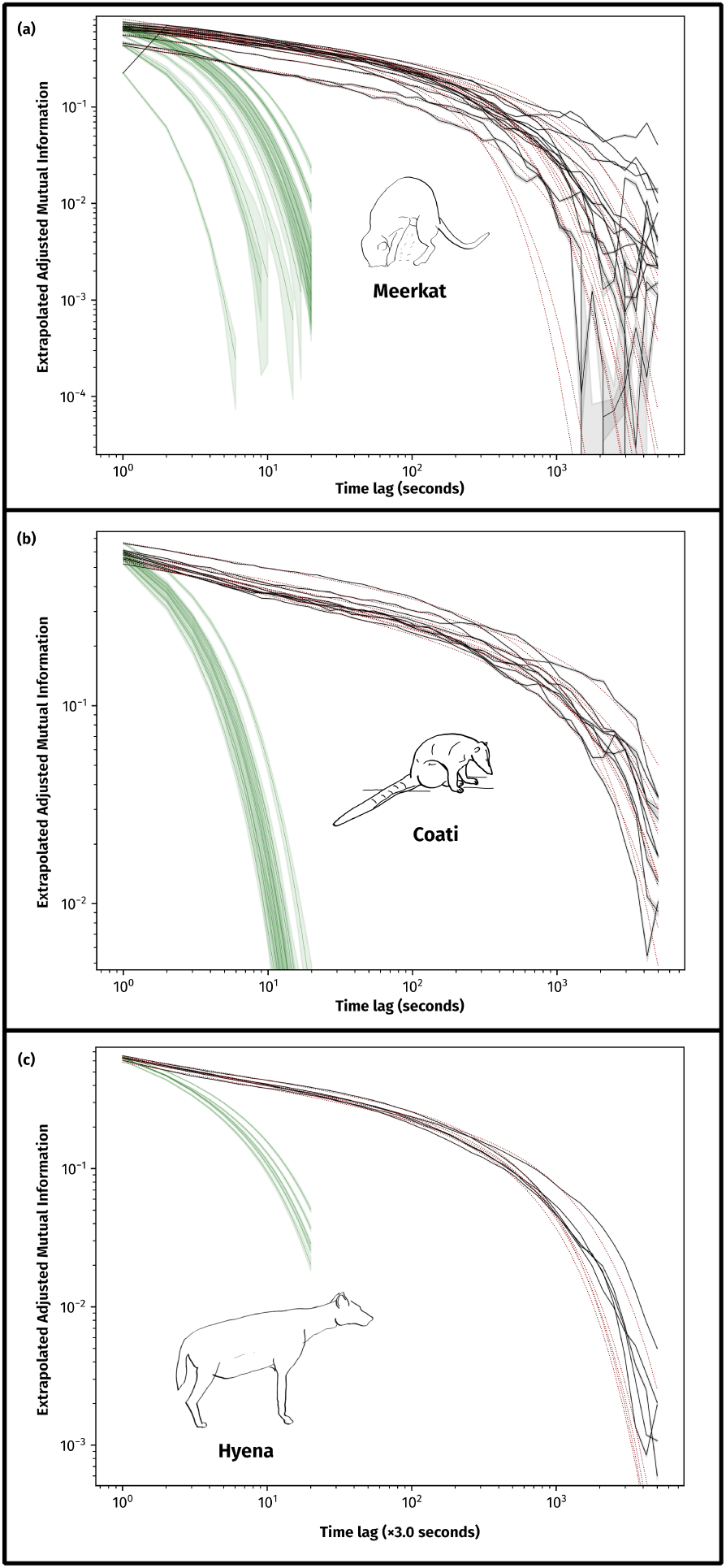
The predictivity of current behavior on future behavior, quantified as adjusted mutual information with finite-size correction (y-axes) between behavioral states of animals at times *t* and *t* + *τ*, (*τ* on x-axes) for (a) meerkats,(b) coatis, and (c) hyenas. Adjusted mutual information (black lines for each individual) always fit to exponentially truncated power-laws (maroon lines). In all species and individuals, behavior remains much more predictive in the future than expected from instant-to-instant dynamics alone (green lines, timelags used for computation are *τ* = 1 … 20). Mutual information tends to increase slightly at high lags for meerkats, leading to worse function fits than in the other two species. When estimates of adjusted mutual information are zero (only observed in meerkats at high lags), black lines intersect the x-axis. This increase might be due to some periodicity in meerkat behavior at these high lags, although the exact origin of this increase is still unknown. Shaded areas indicate 99% confidence intervals.

As a separate way to quantify long-range memory in behavior over time, we also performed Detrended Fluctuation Analyses (DFAs, which quantify long-range memory or ‘dependence’ in time series) [43] for binary sequences of each behavior [as in 6, 23, 44]. The output of a DFA can be interpreted as a power exponent to understand self-similarity and self-affinity in a time-series that is non-stationary (i.e., whose dynamics themselves change over time), and is often used to understand processes with complex self-dependence [e.g., 45, 46]. We found that the exponent *α*_DFA_ was ≈ 1.0 for all behaviors, individuals, and species in our study (see SI Appendix, Table S4 A-C). Such values correspond to so-called ‘1*/f*-noise’, a property attributed to a wide range of complex processes [47, 48]. This result also indicates long-range self-dependence of behavior in these behavioral sequences.

### 2.3 Heavy-tailed bout duration distributions for all behaviors

Several bout duration distributions can lead to decreasing hazard functions. To more fully characterize the variability in bout durations, we quantified bout duration distributions for all behavioral states across all three species, and for each individual separately.

We found that bout duration distributions for a given behavioral state were consistent across individuals (Figure 3), and were most often heavy-tailed (SI Table S2) with almost all individuals showing truncated power-law or lognormal bout duration distributions across all behavioral states. There was no apparent pattern in which states were truncated power-laws and which states were lognormal. The finding of heavy-tailed bout duration distributions implies that it is difficult to describe behavioral bouts with a ‘typical’ time-scale for any of the behaviors quantified here.

**Figure 3:**
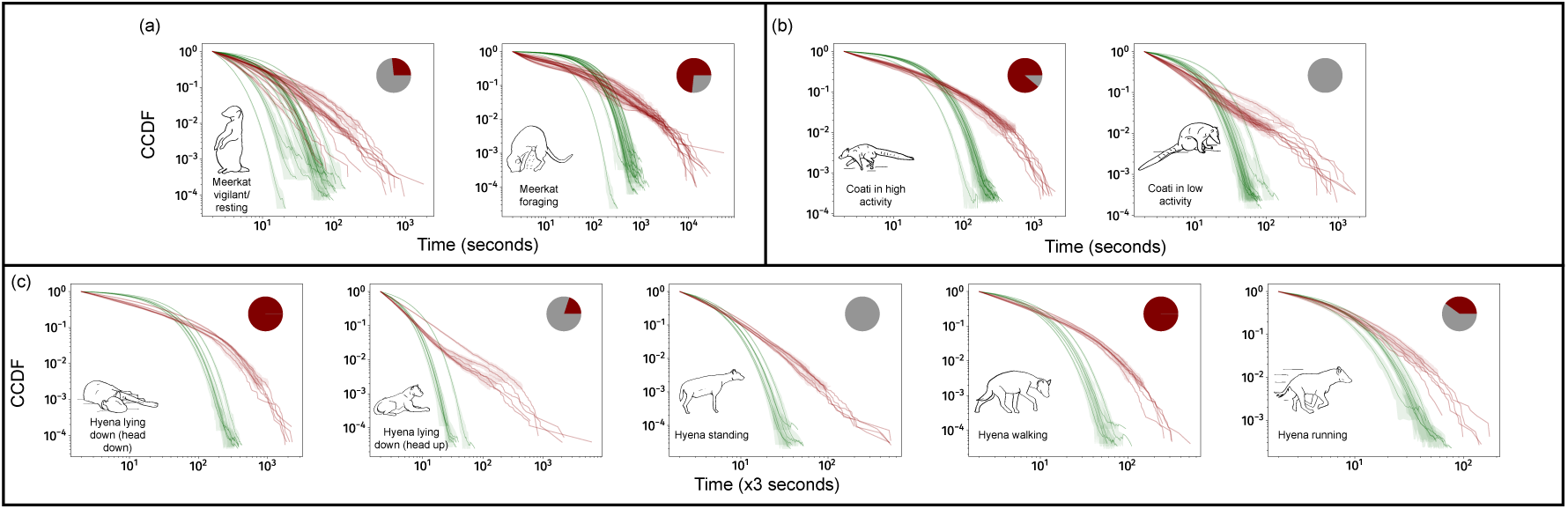
Distributions of bout durations are always heavy-tailed. Panels show observed complementary cumulative distribution functions (CCDFs) for (a) meerkats, (b) coatis, and (c) hyenas. All distributions have a heavy tail, as indicated by the nearly linear (on the log scale) relationship between bout duration and frequency for a significant portion of the distribution. For each behavior, all individuals seem to exhibit very similar bout duration distributions. Each sub-plot represents one behavioral state, and red lines represent bout duration distributions from these states for one individual each. Shaded regions are the 95% CIs of the estimate of CCDFs accounting for finite-size effects. Green lines are best fits for bouts in pseudosequences that preserve instant-to-instant dynamics. As expected, larger bout durations are much more probable in the real data than in these pseudosequences. Inset line diagrams show the behavioral state, while inset pie-charts show the proportion of individuals (in red) that have truncated power-law bout duration distributions (grays are lognormal). No behavior showed an exponential fit for any individual (Table S2 lists best fit distributions). *x*-axes represent duration measured in time-windows (see Methods), and the durations of the time-windows are 1 s for (a) and (b), and 3 s for (c).

### 2.4 Effect of imperfect classification

Errors in the recognition of behaviors can split long bouts into two or more shorter bouts so that even small error rates in behavior identification can have complex, non-trivial effects on the bout duration distributions quantified from their output. In general, we expect this effect to lead to the shortening of longer bouts with higher probabilities as classifier error rates increase, which may lead to false negatives in the quantification of heavy tails. Using simulated behavioral sequences with power-law and exponential bout duration distributions, and classifiers with error rates we could specify (see SI Appendix Text ‘Simulating the effect of classifier error on the quantification of bout duration distributions’, Figure S3A), we found that classifier error indeed produces biases against the detection of heavy tails in bout duration distributions, such that the most error-prone classifiers never detect a heavy tail. Unexpectedly, we also found that, at intermediate error rates, classified behavior sequences with exponential bout duration distributions were characterized erroneously with truncated power-law bout duration distributions (Figure S3B). This spurious detection of heavy tails most likely emerges through the preferential destruction of longer bouts as compared to shorter ones (Figure S3C). Since the longest bouts in our data last up to 10^4^ seconds (3 hours), for error-prone classification as in our simulations to explain our empirical findings, animals would need to exhibit a ‘true’ bout duration distribution with bout lengths two orders of magnitude greater than the longest ones observed in our data (as in Figure S3C), or around 10^6^ seconds (11 days) long. The existence of such long bouts seems exceedingly unlikely, making it highly improbable that classifier error can explain our broad findings of heavy-tailed distributions. Moreover, power-law bout duration distributions were sometimes perceived as lognormal almost independent of the error rate of the classifier. However, exponential bout duration distributions were rarely perceived as lognormal (Figure S3D). It is thus likely that behavioral sequences in the wild have bout duration distributions that have heavier tails than those detected here. Nevertheless, we suggest that future studies investigating the effects of classifier error on behavioral sequence analysis would be important to shed more light on this general issue.

As an additional test of robustness, we re-inferred the behavioral sequences of meerkats using a coarser approach, wherein the behavior labels corresponded to low– and high-activity levels. We redid our analyses on behavioral sequences from this coarser interpretation of meerkat behavioral states (SI Appendix, Text “Repeat analysis on meerkat VeDBA activity levels”). We found that all our results were replicable in these alternative behavioral sequences as well (Fig. S6). This reanalysis serves to rule out the role of classifier artifacts in the observation of long-timescale structure in the data, and also shows that our results are robust to differences in the inference of behavior.

## 3 Discussion

Taken together, our findings highlight common patterns in the organization of behavioral sequences across behaviors, across individuals, and across species. We found that behavioral bouts are described by decreasing hazard functions. We also found a truncated power-law decay in the predictivity of an animal’s present state of its future state, indicating that behavioral sequences are characterized by scale invariance at least up to some long timescale (here roughly 15 minutes for meerkats and coatis, and 45 minutes for hyenas). Further, we found truncated power-law and lognormal distributions of behavioral bout durations, suggesting a multi-timescale organization of behavior, with most behaviors being difficult to characterize by a single ‘typical’ timescale.

Our analyses simplify the behavior of an animal, so that all our behavioral sequences are composed of coarse behavioral labels, and the level of coarseness and temporal resolution are different among study species. Such simplification is useful as it provides great analytical tractability while still retaining temporal patterns of behavioral structure. Still, we note that behavior sequences like these do not describe behavior in its full complexity in an animal, due to which care needs to be taken in inferring and understanding our results. For instance, courtship behavior in a male hyena would form behavioral sequences with short bouts of walking towards and away from a female. That these bouts arise during courtship would tell us a lot about the dynamics of the bouts of movement at this time. Reducing complex behaviors to sequences of simple kinematic behavioral states necessarily loses such contextual information, yet this reduction in complexity also enables a broader perspective on behavioral sequences in general, as we have done here.

The existence of shared structural properties in behavioral sequences suggests that these patterns may be driven by common underlying mechanisms. While these mechanisms remain unknown, we propose here two possible explanations that could give rise to such consistent, widespread patterns. First, positive feedback processes may be a general feature of animals’ behavioral algorithms. Second, the interaction of environmental and/or physiological processes across timescales may introduce memory into behavioral processes that are themselves memory-less.

### 3.1 Behavioral algorithms may contain positive feedback motifs

Our discovery of decreasing hazard functions—where long bouts of behavior tend to get longer— across all behaviors, individuals, and states points towards the possible existence of positive feedback mechanisms in behavioral algorithms. While positive feedback is a common feature in the organization of neural circuits [49–51], these positive feedback mechanisms do not necessarily need to be neurological. Since all three species we investigated here are social, one possible explanation for the convergent patterns is social feedback, where individuals socially reinforce each other’s behavior resulting in prolongation of behavioral bouts. Using simulations of interacting memory-less animals (see SI Appendix, Text ‘Simulating individuals who socially reinforce their behaviors’, Figure S4), we show that such social reinforcement can in principle lead to decreasing hazard rates as observed in our data.

This argument raises the possibility that behavior is broadly governed by positive feedback mechanisms. These positive feedbacks could operate at the level of internal dynamics, social feedback, environmental feedbacks, or a combination of these factors. Such positive feedback mechanisms could also cause the scale invariance of behavioral predictivity, as power-law relations are associated with positive feedbacks [47]. This scale invariance might then break down at longer timescales, as in our results, due to environmental changes occurring at a somewhat slower timescale (e.g., change in time of day, or change in type of habitat as the animal moves), leading to the observed exponential truncations of bout duration distributions seen in many behaviors.

Mechanistically, dynamics such as these could come about if the decision to change behaviors or not is made less and less frequently the longer an animal has been performing a behavior. Similar arguments have been made before in the context of movement decisions to explain Lèvy Flights [52].

The argument that behavioral decision-making is inherently characterized by positive feedbacks has the advantage of being the most straightforward interpretation of our results, in that it attributes the observed patterns directly to the underlying behavioral algorithms at play. However, this argument is difficult to prove or disprove based on behavioral data alone. Future studies which also monitor brain activity might help detect candidate positive feedback motifs in the brain, and shed some light on the validity of this explanation.

### 3.2 Behavioral algorithms may be driven by processes acting at different time-scales

Decreasing hazard functions and the apparently scale invariant decay of behavioral predictivity could also come about as a result of the interaction of several individually memory-less processes, each acting at a different time-scale [5, 20, 53, 54]. Such a combination of processes could arise from different mechanisms, described below.

First, we note that not all mixtures of memory-less processes generate bouts that are described well by all heavy-tailed distributions. Behavior could be conceived as driven by processes at two levels: instantaneous transition rates between behavioral states, and slower dynamics of the transition rates themselves. To explore this simple model, we simulated pairs of exponential distributions with different parameter values. We found that specific combinations of parameters tended to produce data with specific best-fit distributions (SI Appendix ‘Simulating random exponential distribution mixtures that may seem heavy-tailed’ and Figure S5), implying that results like ours are not automatically produced by arbitrary mixtures-of-exponentials. Since it is unlikely that processes acting at only two timescales govern behavior, explanations involving specific, more complex mixtures of a larger (potentially infinite, as in [54]) number of memory-less processes could adequately explain the heavy-tailed distributions observed in our data. For instance, we also showed that in mixtures of three-exponentials with randomly chosen parameter values, around a third produced random numbers that showed truncated power-law best fit (SI Appendix ‘Simulating random exponential distribution mixtures that may seem heavy-tailed’). Explanations following this line of arguments need to explain the potentially shared structuring of mixtures of memory-less processes that generate similar bout duration distributions in all behavioral states across individuals and species. We provide some support for such explanations here.

It has long been argued that animal behavior is hierarchically organized [1, 2], and as such, is probably governed by processes operating at different time-scales at each level of this hierarchy [2, 7, 20, 55]. For example, an animal’s behavior can be coarsely categorized into high and low activity levels. There are bound to be more behavioral states that are encompassed within each of those labels (e.g., standing, sitting, lying down are all low activity behaviors). The rates of state-switching between sub-states within each activity level are likely different from the rate of switching between activity levels. This hierarchy could arguably lead to the perception of prolonged bouts, e.g., of ‘low activity’, while the animal switches between the sub-states encompassed within that label.

Since we expect neither the animal nor the environment to remain the same through time, we expect the same behavior to occur in multiple contexts (e.g., a hyena could be walking in the context of foraging, mate finding, or returning to a den). Different memory-less processes associated with each of these contexts could be sufficient to explain our results, and connect well with the idea of hierarchically organized behavior with different time scales acting at each level [4]. It has moreover been theorized that particular relationships between the timescales relevant to behavior can lead to the appearance of scale invariance [26]. Furthermore, if the animal’s environment varies at a temporal scale slightly slower than the scale at which behavior occurs (which is likely to be true in all animals that move across heterogeneous landscapes), the idea of multiple contexts can be generalized to infinitely many contexts (one for each state of the environment). These contexts could be associated with different (otherwise memory-less) processes, leading to the emergence of heavy tails in bout duration distributions via the combination of a large number of exponential processes. Such a mechanism has recently been theorized and mathematically formalized [53], and could also explain our results.

The observed prevalence of long time-scales in all species considered here can, in these ways, be attributed to multiple intersecting memory-less processes. At the same time, this explanation raises more questions—for instance, how is it that these interacting timescales always combine in such a way to produce apparent scale invariance, particularly with such similar parameter values? Quantifying behavior at multiple time scales requires better analytical approaches [2, 5, 20], as well as further empirical data to test what combinations of times scales may be operating. Further, we need to ask what time scales apply to what contexts, specifically considering the hierarchical organization of behavior: i.e., what time scales apply at what levels of this hierarchy and why.

### 3.3 Possible fitness implications of behavioral sequence structure

The mechanisms hypothesized above do not necessarily imply any kind of adaptive origin. Nevertheless, the possible fitness implications of the structure of behavioral sequences merit consideration.

One aspect of behavioral sequence structure with clear fitness relevance is activity budgeting. Activity budgets are allocations of proportions of total time to different behavioral states [56–58]. The optimal allocation of an activity budget to various behaviors is likely to be driven by the costs and benefits associated with each behavior [59, 60]. Budgets can be adhered to in multiple ways. For instance, an animal may choose to engage in a specific behavior for a set period of time, or may break the behavior into multiple bouts in order to optimize multiple goals such as foraging and predator detection. Algorithms for producing optimal sequences of behavior are not known [but see 61, 62] and it may also be the case that animals use sequences that are ‘good enough’ to meet their needs. It is likely that the large variation that has been observed in bout durations in these data may be attributable to dynamic goals that affect the duration of an engaged behavior, and may somehow be of adaptive value. Future theoretical and empirical studies investigating time budgeting in terms of not only total time allocation but also how such allocations are divided could shed light on this topic.

Another phenomenon to which an adaptive value of scale invariance has been attributed is movement following Lèvy Flights, where step sizes during movement follow a power law distribution [28, 29]. The heavy-tailed distributions of bout durations of movement behaviors we find here accord with prior literature reporting Lévy Flights, however here we have also discovered heavy-tailed bout duration distributions across many behaviors – both those related to movement and those related to rest or inactivity. This raises the intriguing possibility that Lévy flight-like patterns of movement, which have thus far been hypothesized to arise as a consequence of optimal movement strategies [29, 63], could to some extent be explained by general patterns of behavioral dynamics that produce heavy tailed distributions of behavioral bouts [see also 53].

## 4 Conclusions

Analyzing behavioral sequences through three different approaches, we have found remarkable similarities in the dynamics of behavior across three species in the wild. We also found notable similarities across different behaviors in all three species, in contrast to our *a priori* expectations that our analyses should show qualitatively different results for different types of behaviors. These phenomena together strongly suggest that behavior in the wild across species is governed by algorithms with processes likely acting at various time-scales. We theorize that, at least in the species we have studied, these patterns could arise via social or ecological positive feedback mechanisms, via changing environmental conditions, or some combination of both. Independent of their mechanistic explanation, our results imply that decision making processes across our three study species share common structural properties. The surprising cross-species similarity of our results also raises the question of how widespread these phenomena are across the animal kingdom. Future studies quantifying behavioral sequences through biologging and other approaches can expand these analyses to a wide range of species in natural settings, with potential to shed light on general features of behavioral algorithms in animals.

## 5 Methods

### 5.1 Study species and primary data collection

Three species of group-living mammal were considered in this study. Multiple individuals from each species were fitted with collars containing several sensors including accelerometers, which were used to infer behaviors. In each case accelerometer readings were carried out over the course of several days (ranging from 2-47 depending on individual and species; see Table S1). In all cases, accelerometer readings were synchronized with GPS time. Here we give a brief description of each species and the data collection methods. Additional details can be found in SI Appendix, Text ‘Primary data collection and behavioral classifier design’.

Meerkats (*Suricata suricatta*) are social mongooses native to arid parts of southern Africa. They live in highly cohesive groups with a despotic social organization [64]. Meerkats are opportunistic generalists that forage on small invertebrate and some vertebrate prey distributed across their desert habitat by digging in the ground [65]. At night, meerkat groups find shelter in communal burrows, underground structures typically consisting of multiple entrances linked by tunnels, that can persist over many years [66]. Data were collected from all members of three meerkat groups at the Kalahari Research Centre, Northern Cape, South Africa. These individuals wore custom built collars which recorded continuous accelerometery data. Accelerometer data were recorded throughout the day at 10 Hz in some groups, and at 50 Hz in others. Accelerometer readings were synchronized with GPS (Axy-Trek Mini, Technosmart, Colleverde, Italy).

White-nosed coatis (*Nasua narica*) are diurnal members of the procyonid family [67]. Females live in groups with their offspring, whereas adult males are mostly solitary [67–70]. Coatis are omnivorous and spend the majority of the day foraging for leaf litter invertebrates and fruits [71]. Group living coatis groom and play with one another as well as sleep together in tree nests. Data were collected from all 9 members of a coati group in Soberania National Park, Panama. These individuals wore collars that recorded continuous tri-axial accelerometery data at 20 Hz for three hours every day (between 11:00 to 14:00 UTC, corresponding to 06:00 to 09:00 local time), synchronized to GPS data (e-Obs Digital Telemetry, Grünwald, Germany). As continuous accelerometry was limited to three hours a day, we could not measure bouts longer than 3 hours, if any.

Spotted hyenas (*Crocuta crocuta*) are large social carnivores found throughout sub-Saharan Africa. Their bone-crushing jaws and efficient loping gait make them well-adapted for cursorial hunting and scavenging [72, 73]. They live in large, mixed-sex groups where all adults of both sexes are reproductively active and breed non-seasonally [74]. Hyena societies are structured by high degrees of fission-fusion dynamics, where despite living in large groups, they spend most of their time alone or in association with a few group-mates [75]. They are most often active at night and the hours around dawn and dusk, although they can be active at any time of day [76]. Data were collected from five female hyenas residing in the Talek clan in the Maasai Mara National Reserve in southwestern Kenya [77]. These individuals wore Tellus Medium collars (Followit, Sweden) fitted with custom-designed tags (DTAG; Mark Johnson) that collected continuous 24 h tri-axial accelerometer and magnetometer data at 1000 Hz from January 1 until mid-February 2017 [78]. Accelerometry data were down-sampled to 25 Hz for further analysis.

### 5.2 Prediction of behavioral sequences

For all species, available data were divided into discrete non-overlapping time-windows. These intervals were 1 s long for the meerkats and coatis, and 3 s long for the hyenas. Each time-window was labeled with a behavioral category (corresponding to behavioral states), thus predicting behavioral sequences.

With coati accelerometer data, we computed the Vectorial Dynamic Body Acceleration (VeDBA) [79]. VeDBA captures the activity level of an individual. Since VeDBA is bimodally distributed for coatis, we could interpret the two peaks as low– and high-activity states. For each coati, we used a Gaussian Mixture Model with two components to define its activity threshold, and used this threshold to predict its behavioral sequence with the categories low-activity and high-activity.

We predicted more detailed behaviors for meerkats and hyenas using a machine-learning approach. Using previously recorded video data, we generated ground-truth based on a minimal set of behavioral labels corresponding to basic movement states. We then trained random forest classifiers to assign one such movement-state based label to each time-window based on simple kinematic features. Meerkat behavioral labels were foraging, vigilance/resting, and running. Hyena behavioral labels were lying (head down), lying (head up), standing, walking, and running. Additional information about training these classifiers is found in [78] and (see SI Appendix, Text ‘Primary data collection and behavioral classifier design’)

### 5.3 Generation of pseudosequences

As a comparison to the multi-scale dynamics of behavioral sequences observed in our real data, we generated for each individual 30 pseudosequences which preserved the transition probabilities between consecutive states but removed any longer-timescale structure, following ref. [2]. For generating these sequences, we began with the behavioral state at the start of the animal’s true behavioral sequence. For each subsequent moment in time, the next state was then chosen by drawing randomly (with replacement) from all behavioral states that followed a state of that type in the real sequence for that individual. These pseudosequences are thus equivalent to the outputs of first-order Markov models fit to the behavioral sequence of the animal, and serve as null models that exhibit the same finite size effects as the true behavioral sequences.

### 5.4 Quantifying hazard rates

For each individual, we extracted bouts of each behavioral state from predicted behavioral sequences. To avoid extremely small bouts generated by classifier noise, we considered only sequences that were 2 time-windows or longer. For each possible bout length *t*, we counted (a) how many bouts were at least *t* long, and (b) how many bouts were exactly *t* long. Using these, for each *t*, we found the proportion of bouts that had gone on for exactly *t* that stopped within the next time-window. This let us quantify the hazard function, *h*(*t*).

One problem with visualizing hazard functions quantified this way is that the resolution of the hazard function worsens as we move right along the x-axis. For this reason, we have visualized only up to the length of the 100th longest bouts from each behavior although all bouts were used to compute the hazard functions. Our visualizations of hazard functions therefore don’t span the entire range of bout durations accessible to us. However, this step ensures that we have finer resolution throughout our plots of the hazard function than 0.01.

The ‘Running’ behavior of the meerkats occurred too infrequently, and in too short bouts, to successfully visualize their hazard functions.

### 5.5 Quantifying predictivity decay

We quantified how predictive the present behavioral state was of future behavioral states by computing the adjusted mutual information [41] between the time series of behavior and itself lagged by time *τ*. Adjusted mutual information was found using the python package scikit-learn 1.3.1 [80]. 44 integer values of *τ* spaced equally apart on the log-axis between 1 and 5000 were chosen, and for each *τ* adjusted mutual information between states at time *t* and *t* + *τ* were computed.

Mutual information does not obey the Central Limit Theorem, and a finite-size correction needed to be performed to calculate the value of the metric as data size gets very large [42]. For this finite-size correction, we followed a modified version of the method used in ref. [42]. We computed the adjusted mutual information for different subsets of the data, and extrapolated the adjusted mutual information as dataset size approached ∞. For a dataset with *N* behavioral state pairs (behavioral states at *t* and *t* + *τ*), we drew five subsets for each of five sizes *n* ∈ {0.5*N*, …, 0.9*N* }. We then performed a linear regression between 1*/n* and adjusted mutual information values, and extracted the value of the *y*-intercept (1*/n* → 0) as the extrapolated ‘true’ value of the desired adjusted mutual information.

Finally, to compute how behavioral predictivity decays, based on these adjusted mutual information values, we fit functions of *τ* and computed *R*^2^ values for goodness-of-fit comparisons. The functions we fit were an exponential decay (*me^−λτ^*), a power-law decay (*mτ ^−α^*), and a power-law decay with exponential truncation (*mτ ^−α^e^−λτ^*). This was done separately for each individual.

### 5.6 Detrended Fluctuation Analyses

We performed Detrended Fluctuation Analyses (DFAs) as a separate way to show long-range self-dependence in behavioral sequences. This is a standard analysis for dealing with non-stationary processes to understand how fluctuations in time-series depend on temporal scale. The outputs of DFA analyses (*α*_DFA_) can be used to make inferences about this scaling.

For each behavior, we generated binary time series (1 when the animal was in the focal behavior, –1 when it was not) for each behavior. We then performed DFA using the python package nolds [81], and recorded values of the DFA exponent (*α*_DFA_)for each behavioral state and each individual.

### 5.7 Fitting and comparing distributions

As with hazard rates, we extracted bouts of each state for each individual and excluded the 1 time-window bouts. We then fit these bouts to discrete exponential, lognormal, power-law, and truncated power-law distributions. For fitting, we followed recent statistical developments [82], and used the library powerlaw [83] (v1.5). For each distribution *A*, we calculated the log-likelihood

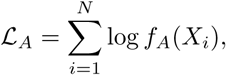

where *X_i_*are the observed bout durations, *f_A_*is the estimated maximum likelihood probability density function for distribution *A*, and *N* is the total number of data-points. Using L*_A_*, we then computed the Akaike Information Criterion [84] as

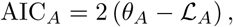

where *θ_A_* is the number of parameters in the distribution *A*. The distribution with the lowest AIC was chosen as the best-fit distribution.

We only fit distributions if at least 250 bouts of a behavior could be found for an individual. This condition was satisfied by all behaviors in all individuals across all species except the ‘Running’ state in meerkats, where we could not quantify bout duration distributions with the available number of bouts.

### 5.8 Code and data availability

All code used to analyze behavioral sequences and perform simulations in this study is publicly available as a GitHub repository [85], as is all code used to infer behavior for meerkats [86], coatis [87], and hyenas [88]. In this initial submission, data are made available to reviewers at this URL to access and download: https://owncloud.gwdg.de/index.php/s/7aejioFxFBIlVBU. On acceptance, code used will be assigned a permanent DOI using Zenodo, and all data used here will be made accessible to the public using Dryad.

### 5.9 Ethics

The data used in this study were collected in accordance with institutional ethical guidelines and with approval from relevant local authorities. The collection of data on meerkats was approved by ethical committees at the University of Pretoria, South Africa (permit: EC031-17) and the Northern Cape Department of Environment and Nature Conservation (permit: FAUNA 1020/2016). Research on coatis was approved by the Smithsonian Tropical Research Institute (STRI) Animal Care and Use Committee (IACUC Assurance number 2017-0815-2020). Hyena field methods were approved by Kenya Wildlife Service under permit KWS/BRM/5001 to KEH, and were also approved by the IACUC at Michigan State University under approval PROTO201900126.

## Acknowledgements

We thank Baptiste Averly, Frants H Jensen, Marta B Manser, and other members of the Communication & Coordination Across Scales project team for contributing to the data collection underlying this work, and for helpful discussions. We also thank Alison Ashbury, Antonio Costa, and Hester Brønnvik for feedback on earlier versions of the manuscript. ASP, BTH, KEH, and MAR acknowledge funding from Human Frontier Science Program (HFSP) Research Grant RGP0051/2019. PM acknowledges support from a Deutscher Akademischer Austauschdienst (DAAD) fellowship. ASP acknowledges funding from the Gips-Schüle Stiftung and the Max Planck Society. The work was additionally funded by the Deutsche Forschungsgemeinschaft (DFG, German Research Foundation) under Germany’s Excellence Strategy – EXC 2117 – 422037984. MCC acknowledges support from a Packard Foundation Fellowship (grant no. 2016-65130), a grant from the National Science Foundation (grant no. NSF BCS 1514174), and by the Alexander von Humboldt Professorship endowed by the Federal Ministry of Education and Research. The long-term study of meerkats is currently supported by the MAVA foundation, funding from the European Research Council (ERC) under the European Union’s Horizon 2020 research and innovation program (No. 742808 and No. 294494), and a grant from the Natural Environment Research Council (Grant NE/G006822/1) to Tim Clutton-Brock, as well as by grants from the University of Zurich to Marta B. Manser. Parts of the meerkat study were done when VD was funded by Minerva Stiftung and Alexander von Humboldt Stiftung postdoctoral fellowships. We thank the Republic of Panama for permission to conduct the coati study. We thank STRI for their assistance and use of their research facilities. We thank Patrick Paetzold for coati collar construction assistance. For help with coati fieldwork and data collection, we thank Carolina Mitre Ramos, Brandol Ortega, and Lucía Torrez. We thank Rachel Page and Melissa Cano for their support during fieldwork. Part of the funding for the coati study was provided by the Alexander von Humboldt Professorship, endowed by the Federal Ministry of Education and Research awarded to MCC. The long-term study of spotted hyenas was supported by U.S. National Science Foundation (NSF) Grants OISE1853934 and IOS1755089 to KEH. We thank the Kenyan National Commission for Science, Technology and Innovation, the Narok County Government, the Mara Conservancy, the Kenya Wildlife Service, Naboisho Conservancy, and the Wildlife Research Training Institute for the permission to conduct the hyena research. We also thank all those who assisted with hyena data collection in the field, especially Malit Ole Pioon. We thank Rebecca LaFleur, Kenna Lehmann, Morgan Lucot, Max Martin, and Benson Pion for their help in hyena fieldwork and data curation, and Mark P Johnson for providing the DTAG technology used in hyena collars. We thank the two reviewers for their thorough comments, which have led to significant improvements in the robustness and rigor of our results and interpretations.

## Author contributions

The study was conceptualized by PM and supervised by ASP. VD, LJU, and ASP collected the data on meerkats; EMG, MCC, and BTH collected the data on coatis; and KEH, ASG, and ASP collected the data on hyenas. The meerkat data were collected in cooperation with the Kalahari Meerkat Project, headed by Marta B. Manser, and the hyena data were collected in cooperation with the Mara Hyena Project, headed by KEH. AN, EMG and ASG contributed to labeling of ground truth video data in meerkats, coatis, and hyenas respectively. PM designed systems to infer behavior from accelerometer data, and designed and performed all analyses presented. KB and ASP performed a thorough code review. PM wrote the first draft of the paper with inputs from EMG, VD, and EDS. All authors contributed to and approved the final draft.

