## Supporting Information for "Behavioral sequences across multiple animal species in the wild share common structural features"

February 20, 2025

**6 Repeat analysis on meerkat VeDBA activity levels** **11**

#### 1 Primary data collection and behavioral classifier design

In this section, we describe the biology and behavior of all three study species and how we inferred each of their behavioral states using continuous accelerometer recording. Because the tools used differ among the different study species, we explain the specifics of each of them below separately. However, common to all methods of analysis is the Vectorial Dynamic Body Acceleration (VeDBA) metric calculated for each time point, which is indicative of the level of activity of the animal in that instant [1].

We used VeDBA and other features to arrive at a classification system for each species that associated a behavioral state with each point of data available for each individual. In all cases, labels corresponding to behavioral states are attributed to time-windows (1 s or 3 s long) based on accelerometer data. Behaviors inferred this way constitute long behavioral sequences for each individual, providing rich insights into the behavioral dynamics of these animals Table 1.

#### 1.1 Meerkats

##### Behavior and biology

Meerkats (*Suricata suricatta*) are social mongooses native to arid parts of southern Africa. They live in highly cohesive groups with a despotic social organization [2] of a dominant pair monopolizing most of the breeding opportunities [3], and much weaker social hierarchies among subordinate group members [4]. During the day, meerkat groups move as a cohesive unit through their territory while foraging by probing and digging in the soil for prey [5, 6]. When foraging between vegetation, meerkats have a limited view of their surroundings, and rely heavily on acoustic communication for monitoring the positions of their group mates [6, 7]. Meerkats frequently show sentinel behavior where one individual climbs to an elevated position and scans the area for the presence of predators. During foraging, meerkats frequently encounter aerial and terrestrial predators (on average they seek shelter every 40 min [8]), which makes a vigilance state very important. At night meerkat groups shelter in communal burrows, underground structures typically consisting of multiple entrances linked by tunnels, that can persist over many years [9].

We collected the data for this study at the Kalahari Research Centre, Northern Cape, South Africa in May-August of 2021. The individually marked meerkat population on site is fully habituated to human presence including behavioral observations from $< 1$ m. Meerkat groups on site are continuously monitored and long term individual and behavioral data is collected by a team of trained researchers. All procedures were approved by ethical committees of the University of Pretoria, South Africa and the Northern Cape Department of Environment and Nature Conservation.

We deployed custom built collars in four meerkat groups of 12-15 individuals. Tracking collars consisted of a GPS and ACC unit (Axy-Trek Mini, Technosmart, Colleverde, Italy) with a miniature audio recorder (Edic-mini Tiny+ A77), mounted on a 5-millimeter-wide leather strap and weather sealed by 2-part epoxy glue. The audio unit was positioned below the chin of the animal and the GPS unit and antenna were positioned at the back of its head for improved reception. Assembled tags weighed 22 - 24g, well below the accepted threshold of 5% body weight of an adult animal.

We programmed the GPS and audio loggers to record for 3 h each day during times when meerkats typically forage within their territory while moving as a group (either in the morning after the group had left the sleeping burrow, or in the afternoon before returning to it), over the course of 5–13 days per group. During recording sessions, an observer noted the times of any group-level disturbances (predator alarms, inter-group encounters, and resting periods) [10]. Accelerometer loggers were programmed for continuous sampling at 50 Hz or 10 Hz depending on the group, with 10 bit resolution and 8g acceleration range.

**Table 1:** Data availability for (a) meerkats, (b) coatis, and (c) hyenas. Each row represents a separate individual. The columns with state behaviors as headers present the number of bouts of those states that were available.

(a)

| Data available (hours) | Days collared | Foraging | Running | Vigilance / resting |
|---|---|---|---|---|
| 167.024 | 8 | 5495 | 99 | 5411 |
| 46.543 | 4 | 3118 | 23 | 3102 |
| 71.376 | 5 | 3715 | 43 | 3678 |
| 27.608 | 2 | 1375 | 17 | 1359 |
| 167.031 | 8 | 7333 | 114 | 7227 |
| 68.213 | 4 | 3011 | 65 | 2948 |
| 122.239 | 6 | 4970 | 121 | 4853 |
| 97.744 | 5 | 5260 | 44 | 5218 |
| 319.001 | 15 | 16519 | 101 | 16426 |
| 318.981 | 15 | 12584 | 195 | 12399 |
| 49.478 | 3 | 1855 | 25 | 1831 |
| 312.628 | 9 | 27683 | 110 | 27107 |
| 169.097 | 8 | 7885 | 71 | 7818 |
| 306.542 | 16 | 9949 | 144 | 9813 |
| 298.817 | 15 | 16109 | 198 | 15919 |

(b)

| Data available (hours) | Days collared | High activity | Low activity |
|---|---|---|---|
| 45.742 | 16 | 5206 | 5205 |
| 45.509 | 16 | 4728 | 4727 |
| 46.346 | 16 | 5159 | 5157 |
| 20.556 | 7 | 2372 | 2370 |
| 46.168 | 16 | 5150 | 5147 |
| 45.833 | 16 | 5671 | 5666 |
| 46.638 | 16 | 7914 | 7908 |
| 43.362 | 15 | 4586 | 4582 |
| 45.694 | 16 | 4978 | 4976 |

(c)

| Data available (hours) | Days collared | Running | Lying (head down) | Lying (head up) | Standing | Walking |
|---|---|---|---|---|---|---|
| 334.501 | 43 | 4998 | 24074 | 37770 | 54611 | 34758 |
| 364.747 | 47 | 5183 | 21463 | 54787 | 71737 | 40005 |
| 99.944 | 14 | 1214 | 7626 | 13838 | 18692 | 10571 |
| 349.029 | 45 | 3375 | 19043 | 39753 | 54948 | 32665 |
| 343.114 | 44 | 3758 | 21930 | 49892 | 57619 | 29641 |

##### Behavior classification

Behaviors were manually labeled for each video using BORIS [11], a software commonly used for video annotations. Alongside VeDBA, for every 1 s non-overlapping time-window, we calculated the mean, variance, minimum, and maximum of readings along each of the x, y, and z accelerometer axes. These values were used as features to train a random forest classifier against the behaviors from video annotations. For simplicity, we chose a 3-behavior ethogram with the behavioral states vigilance, foraging, and running, which is representative of meerkat behavior during their foraging trips into their territory during the day.

In all testing methods, we evaluated the accuracy score (fraction of behaviors correctly identified), $F_1$ score (weighted average of harmonic means of precision-recall values), and confusion matrix (fractions of each behavior $i$ identified by the classifier as each behavior $j$ for all behavior pairs). The classifier was first evaluated using randomized testing, wherein after shuffling all available data, 85% was used for testing. After this, to test generalizability, we used a leave-one-individual-out approach, wherein the classifier was tested on each individual after training with data from all other individuals.

We found that the classifier performed well, with an accuracy of 95.2% ($F_1 = 0.94$) under randomized testing and 93.6% ($F_1 = 0.93$) under individual-wise testing, suggesting that the classifier generalizes well to new individuals. While vigilance and foraging are well inferred, instances of meerkats running were not well identified by the classifier Figure 1. This is most likely because in meerkats running is a rare behavior occurring in short bursts: In 1 h and 37 min of available training data from video annotation, only 14 s (6 bouts from across 3 individuals) of running data was available. This might explain why the running state is not identified well when comparing across individuals. Nevertheless, the classifier is quite useful in distinguishing the more common vigilance and foraging states.

#### 1.2 Coatis

##### Behavior and biology

White-nosed coatis (*Nasua narica*, hereafter 'coatis'.) are a highly social species in the family Procyonidae. Females of all ages, and males under two years of age live in groups. Adult males tend to be solitary, joining female groups during the mating season [12, 13]. Group sizes range from 6 to over 30 individuals [14, 15]; and these coatis typically closely related to each other [16]. A range of behaviors have been observed in coatis, from aggression and mobbing, to grooming and playing [13, 14, 17, 18]. Coatis spend the majority of the day foraging on the ground for invertebrates found in the leaf litter and soil. Coatis also spend a significant amount of time foraging in and below fruiting trees. During the night, coatis will typically sleep together in tree nests to minimise their risk of predation [13]. During a typical day, coatis have a mid-day rest period [13]. Females leave the group to give birth and return to the group when the young are 5-6 weeks old [13]. This study was conducted before and during the period when the adult female left the group to give birth. In this study, we collected collar data from all individuals from one group that consisted of one adult female, two sub-adult males and 6 juveniles. Each group member wore a custom built collar which housed a GPS and accelerometer unit which was positioned at the back of their head (e-Obs Digital Telemetry, Grünwald, Germany), and two audio recorders were positioned under their neck (Soroka 18E, TS-Market). GPS and audio recorders were mounted on a 12-millimeter-wide strap

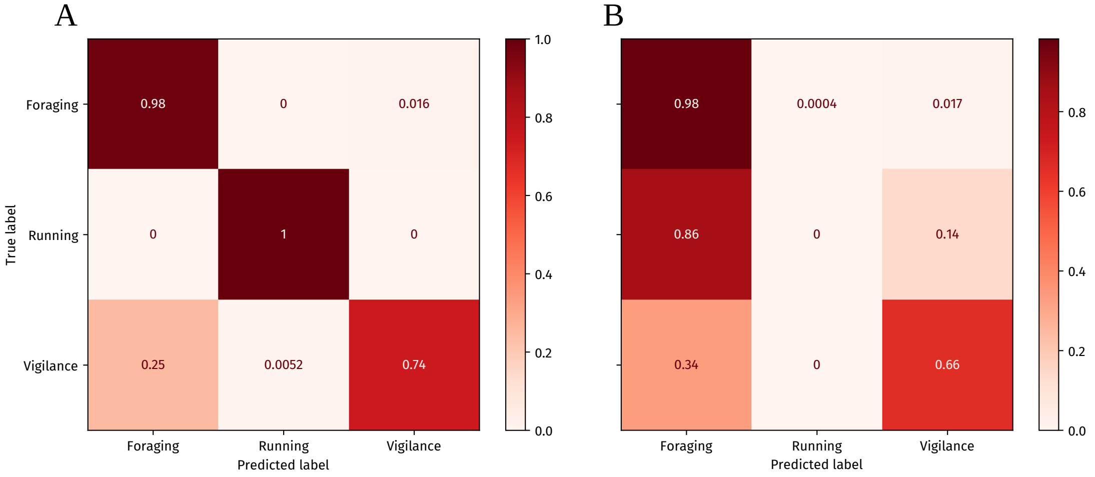

**Figure 1:** Confusion matrices for the meerkat behavioral classifier with (A) randomized testing with 15% of the data, and (B) individual-wise testing. The classifier generally performs well. The 'running' behavioral state is hard to identify across individuals.

and weather sealed using a 3D-printed case and 2-part epoxy glue. Collars weighed less than 5% of the animals body weight. Accelerometer loggers had a 12 bit resolution in each axis with a measurement maximum of 2g acceleration. All collars were programmed to record 1 Hz GPS and 20 Hz continuous accelerometer data from 6 AM to 9 AM local time for 16 days. From 9 AM to 6 AM the next day, 6.9 s accelerometer recording bursts at 20 Hz were recorded once per minute. From 9 AM to 6 PM, 6 s GPS bursts were recorded once every 10 minutes. From 6 PM to 6 AM, 6 s GPS bursts were recorded once an hour. Audio recorders were programmed to record from 6 AM to 9 AM at 24 kHz.

##### Behavior classification

Behaviors were inferred for the continuous 20 Hz recording intervals—between 6 AM and 9 AM everyday. After dividing available accelerometer data into non-overlapping 1 s time-windows, VeDBA values were calculated during the continuous recording interval from 6 AM to 9 AM. We found that VeDBA was distributed in a bimodal manner for each individual coati (Figure 2), which made it convenient to choose low- and high-activity states corresponding to each peak in the distribution. We fit a Gaussian Mixture Model with two components to log VeDBA, and used this model to assign low-activity and high-activity labels to generate coati behavioral sequences for each individual. Since Gaussian Mixture Models were fit for each individual separately, the definitions of low- and high-activity are individual specific, and account for the individual's overall tendency to be active or inactive.

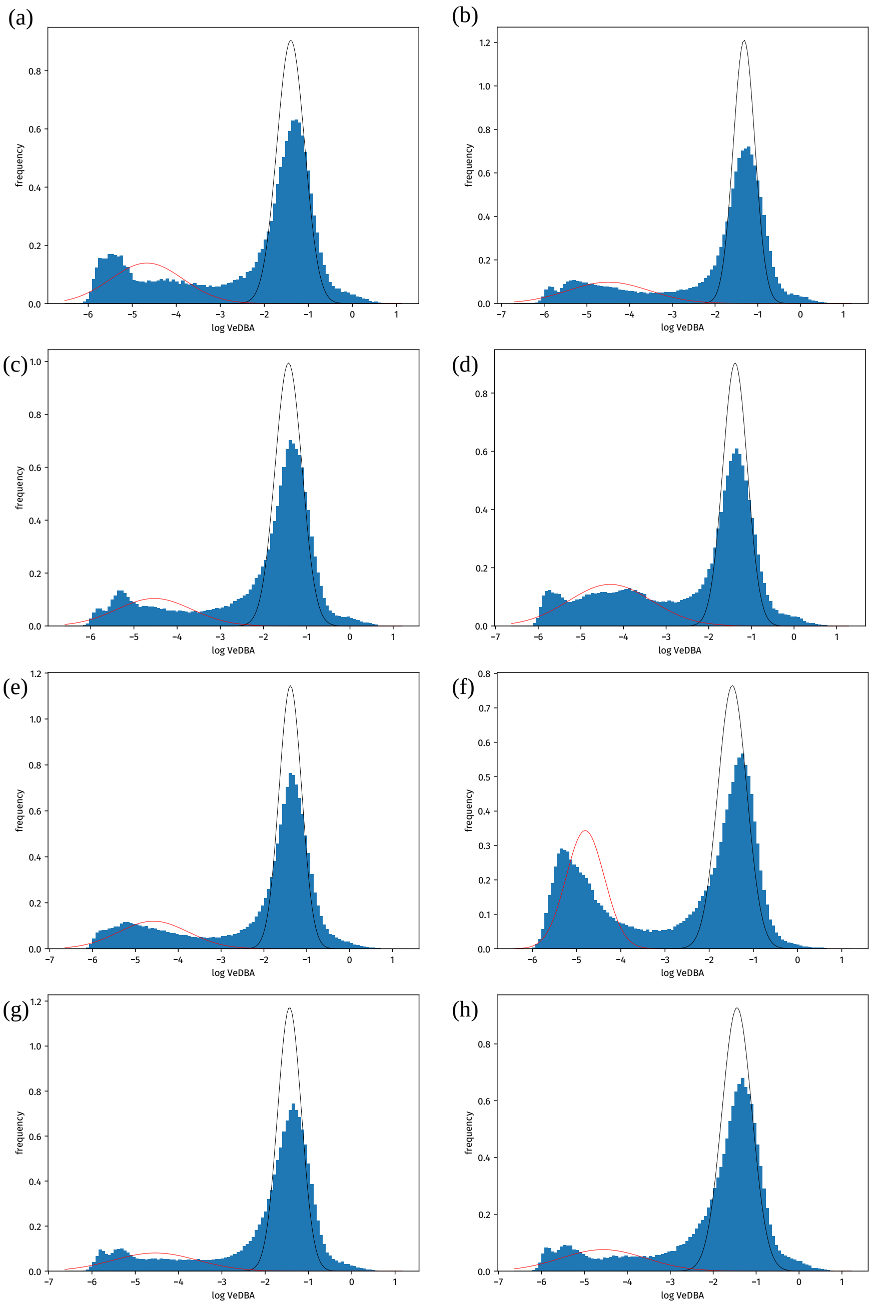

**Figure 2:** Distributions of VeDBA for 8 coatis are shown here (all 9 followed the same bimodal distribution). Normal distributions predicted for the two components (low- and high-activity) by our Gaussian Mixture Model are plotted in red and black.

##### 1.3 Hyenas

###### Behavior and biology

Spotted hyenas are the most social and well-studied of the four members of the Hyenidae family [19, 20]. They live in large groups structured by high-degrees of fission-fusion dynamics, where individuals associate in subgroups of one or more individuals that change composition fluidly throughout the day [21]. Group-mates separated by long distances can communicate through long calls called whoops, which can sometimes function to recruit large numbers of group-mates for cooperative behaviors like competition with sympatric carnivores or other groups of hyenas [22, 23]. Although they can be active at any time of day or night, spotted hyenas are most active at night and in the hours around dawn and dusk [24]. In this population, hyenas forage primarily by hunting and killing large ungulates, and do so both in groups and alone [25]. Overall, because of the propensity for hyenas to act alone or with group-mates, behavioral bouts in spotted hyenas are strongly influenced by both individual motivation and the influence of social partners.

In this study, we deployed multi-sensor collars on five adult female spotted hyenas in the Talek clan in the Maasai Mara National Game Reserve in southwestern Kenya. Collars were Tellus Medium units made by Followit Sweden AB, fitted with an additional custom-designed tag (DTAG 4; Mark Johnson) that recorded continuous tri-axial acclerometer and magnetometer data at 1000 Hz. To train the behavioral classifier, video data was collected from collared hyenas during 12 playback experiments where recruitment whoops [22] were played to resting hyenas from a speaker. Responses to the playback stimulus were recorded from a nearby vehicle, and behavioral states were annotated from this video using labeling tools developed by Mark Johnson and Frants Jensen. Hyenas were coded as being in one of five mutually exclusive behavioral states representing a spectrum of physical movement states: lying down with head lowered, lying down with head raised, standing, walking, and loping (a cursorial gait).

###### Behavior classification

Video data were manually annotated. Using the available triaxial accelerometer data, we computed VeDBA values and the mean, variance, minimum, and maximum, each for x, y, and z accelerometer axes, for every non-overlapping 3 s time-window of data, for each hyena. We then trained a random forest classifier to distinguish 5 hyena behaviors: lying down, lying down with an upraised head, standing, walking, and loping. The classifier performed well, with 92% accuracy ($F_1 = 0.91$) using randomized testing, and accurately predicted the hyenas' typical daily activity patterns. The validation and uses of this classifier have been presented in ref. [26].

#### 2 Simulating the effect of classifier error on the quantification of bout duration distributions

When bouts of behaviors are interpreted by an error-prone classifier, long bouts are likely to be broken by erroneous classifications into smaller bouts. Longer true bouts are more likely to contain multiple errors, and are thus split into several more bouts. The distribution of the bouts perceived after classification is thus non-trivially dependent on the true bout duration distribution.

We simulated several behavioral sequences with both heavy-tailed power law bout duration distributions as well as memory-less exponential bout duration distributions. We then interpreted these sequences through simulated classifiers of different error rates. Fitting candidate distributions to the resulting perceived behavioral sequences, we quantified the likelihood of an error-prone classifier to correctly or incorrectly detect a heavy-tailed distribution as a power-law or truncated power-law. We also found the likelihood of lognormals being detected in both these cases.

#### 2.1 Simulation details and results

Consider an agent exhibiting a behavioral sequence with behaviors 'A' and 'B'. We measured a simulated feature, $\gamma$. The value of $\gamma$ was drawn at each second from a distribution with PDF $f_A$ or $f_B$ depending on the behavior of the agent at that time. For both distributions, we chose normal distributions with $\sigma^2 = 1$, and chose the means $\mu_A$ and $\mu_B$ to obtain a desired error rate in a Bayes Classifier, which provides the maximum accuracy possible from the data [27].

By construction, we chose $\mu_B = -\mu_A$. For a desired error rate, $l$, we chose

$$\mu_A = \Phi^{-1}(l),$$

where $\Phi$ is the Cumulative Distribution Function (CDF) of the standard normal distribution, and $\Phi^{-1}$ is its inverse (also called the percentile function).

Since the means were equidistant on both sides of zero, an optimal (Bayes) classifier decides that all values of the feature, when they are less than 0, correspond to the state 'A', and when they are more than 0, correspond to the state 'B'. When $\mu_A$ and $\mu_B$ are set closer to 0, the predicted behavioral sequence has a higher error rate.

The true behavioral sequence was created by drawing bouts from desired bout duration distributions. We performed two tests. In the first test, we drew bout durations from an exponential distribution with parameter $\lambda = 0.01$ for both states. In the second test, the bouts of both states were drawn from a power-law distribution with parameter $\alpha = 1.5$. Behavior sequences were generated by drawing bouts of alternating behaviors until we had 1000 bouts of each behavior, or until the sequence was 100,000 seconds long, whichever occurred first.

20 Bayes classifiers were chosen with error rates from 0.0025 (highly accurate) to 0.5 (randomly guessing classifier), and were used to generate predicted behavioral sequences. Candidate distributions were fit to these sequences as described in the main text. This was repeated 500 times each for exponential and power-law distributed bouts. For each classifier, we then checked how many times scale invariant distributions (power-law or truncated power-law) were detected, and how many times lognormal distributions were detected.

We found that true power-laws are, overall, lost with increasing classifier error. Interestingly, memory-less processes were extremely likely to be mistaken for processes with heavy-tailed bouts at intermediate classifier errors (Figure 3 B). However, this effect is accompanied by an accelerated loss of long bouts, leading to a distribution which appears somewhat like a truncated power-law (Figure 3 C) with a less heavy tail than the exponential distributions in the true behavioral sequence. We also found that lognormal distributions could be detected in perceived behavioral sequences when the true bout duration distributions were power-laws. Conversely, the incorrect perception of lognormal bout duration distributions when the true bout duration distributions were exponentially distributed is very rare.

While the misdetection of bout duration distributions due to the loss of long bouts to error-prone classification could contribute to us finding heavy-tailed bout duration distributions in our three species, the mechanism behind this effect makes it very unlikely. We see several very long bouts of up to $10^4$ seconds in our data. For our heavy tails to be caused by this effect of intermediate errors implies that, in truth, there are several more such long (or much longer) bouts of all behaviors that have been cut up by the intermediately error-prone classifiers. The existence of such long bouts more than what we have found is doubtful.

Future studies can focus on quantifying this effect in agents showing non-identical behaviors, and eventually in agents showing more than two behaviors.

### 3 Simulating individuals who socially reinforce their behaviors

Decreasing hazard rates, associated with reinforcement in some form, are seen whenever longer bouts come about in contexts with low bout switching probabilities. An illustration of this is the case of social reinforcement of behaviors. Consider a group of animals that show two behaviors. Suppose that, at each point of time, each animal has some probability of switching from its current behavior. Suppose also that the probability increases number of its group members performing the other behavior. In such a situation, we expect longer bouts of a given behavior to be associated with times when most of the group was performing that behavior, and therefore associated with a low instantaneous probability of the behavior ending. This social reinforcement must present itself as a decreasing hazard function.

#### 3.1 Simulation details and results

We simulated groups of 5, 25, and 125 individuals who could perform two behaviors, 'A' and 'B'. Every step, the probability of an individual exiting its current state is based on the proportion of individuals in the other state. In this specific case, for an animal in behavior 'A', we set the probability of switching to 0.01 when none of the animals are doing "B", 0.1 when all animals are doing 'B', and scaled exponentially between these values when some of the individuals are doing 'B' (i.e., $\Pr(A \to B) = 10^{-1-p}$ where $p$ is the proportion of individuals doing 'A'). This rule is symmetrically applied to animals doing 'B' as well ($\Pr(B \to A) = 10^{p-2}$). These groups were randomly initialized and simulated for 50000 units of time. Such simulations were run ten times for each group size. Hazard functions of behavioral bouts were computed for each individual as in the main text.

We found a decreasing hazard function for bout durations of these behaviors in all three group sizes(Figure 4). Such a decreasing functions are seen because, in an individual, long bouts of a behavior are likely to arise from situations when most or all of the group is performing the same behavior. A simple but incorrect interpretation would be that the individual progressively becomes less and less likely to exit a behavior the longer it has done it. As this simulation shows, different contexts being associated with longer and shorter behaviors are sufficient to induce memory in memoryless systems, computed here as decreasing hazard functions. We note also that the framework set up here is useful beyond this study: future studies might explore the nature of hazard functions and other metrics on such socially reinforcing groups of behaving agents, and ask how exactly transition probabilities, heterogeneity, and the number of possible behavioral states might influence the behavioral dynamics of social animals.

### 4 Simulating exponential distribution mixtures that may seem heavy-tailed

Certain mixtures of exponential distributions can produce data that appear to be distributed with a heavy tail [28]. It was, however, not clear how easily arbitrary mixtures of exponentially distributed data appear heavy-tailed, and which mixtures fit well to which heavy-tailed distributions. We explored the parameter space of mixtures of two exponential distributions and found the best-fit distribution on data generated by these mixtures.

To generate the mixtures, we followed the following steps: We started with an exponential distribution with parameter $\lambda_1 = 10^{-2}$, corresponding to an average bout length of 100 s. We then chose another exponential distribution, whose parameter $\lambda_2$ was varied from $10^{-6}$ to $10^{1}$ corresponding to a bout duration between 100,0000 s and 0.1 s respectively. Finally, another parameter, $p$, was varied between 0 and 1 at uniform intervals of 0.01. $p$ determined the mixing proportions of the two distributions. Random numbers were then generated following the probability distribution function

$$f(x) = p\lambda_1 e^{-\lambda_1 x} + (1-p)\lambda_2 e^{-\lambda_2 x}.$$

100 such numbers were drawn from each mixture. To fit the candidate distribution to these numbers, we followed the same method as in the main text.

The exploration of mixtures of more than two parameters this way is hampered by a curse-of-dimensionality problem. For a mixture of $n$ exponential distribution, we need to specify $2n-1$ parameters ($n$ exponential distributions, and $n-1$ weights). However, we begin this exploration here by creating 35,000 random mixtures of 3 exponential distributions each. From each mixture, we draw 1000 numbers, and perform distribution fitting as described in the main text. To generate a mixture of exponentials, three $\lambda_i$ values were drawn so that $\lambda_i \sim \mathrm{Unif}([0,1])$. We drew the weights such that $(p_1, p_2, p_3) \sim \mathrm{Dirichlet}(1,1,1)$. As before, we initialized the mixture with the density function

$$f(x) = \sum_i p_i \lambda_i e^{-\lambda_i x}$$

#### 4.1 Simulation results

In the two-exponential mixtures, only specific regions in parameter-space generated data that had truncated power-law or lognormal best fits (Figure 5). This means that, if behavior is governed by processes acting at different timescales, particular timescales must operate to create bout duration distributions like those seen in our results (Main text). Although one might expect an anti-symmetry about the $\lambda_1 = \lambda_2$ line, this does not happen. This is likely a result of our distributions producing discrete-valued outputs, and our protocol's exclusion of very short bouts from the best fit.

In three-exponential mixtures, while we did not attempt any formal parameter-space exploration, we found that 33.6% of all random mixtures had truncated power-law best fits (35.7% lognormal; 30.6% exponential). Similar analyses with three or more exponential distributions in the mix, as well as a better mathematical formalization and exploration of this problem, can shed light on general properties of mixtures of memoryless processes that lead to apparently heavy-tailed bout duration distributions. This will also be useful in understanding the multi-scale nature of behavioral dynamics.

### 5 Additional analyses and tables

#### 5.1 Bout duration distribution fits

All behaviors were exhibited in bouts whose distributions followed lognormal or truncated power-law distributions Table 2. These fits have been determined using Akaike Information Critera (see Materials and Methods in main text).

#### 5.2 Predictivity decay

As described in the main text, we computed how the mutual information between each animal's behaviors at time $t$ and $t + \tau$ decayed with $\tau$, i.e., how the predictivity of an animal's current behavior decayed into the future. We found that mutual information always decayed as a truncated power-law, with very similar power-law exponents $\alpha$ across species, across individuals, and across behaviors (Table 3).

#### 5.3 Detrended Fluctuation Analyses

We found that all individuals, across species, showed long-range dependence ($\alpha_{\mathrm{DFA}} > 0.5$) and in fact $\alpha_{\mathrm{DFA}} \approx 1.0$ for all behaviors (Table 4).

### 6 Repeat analysis on meerkat VeDBA activity levels

As a further test against classifier artifacts, we re-inferred meerkat behavioral sequences with the unsupervised clustering of VeDBA approach used for the coatis (section 1). Additionally, we chose 2 s (instead of the 1 s used in the analysis in the main text) as the length of the time-window for this analysis, to illustrate the robustness of our test to different choices of window length.

Meerkat VeDBA were calculated with a 2 s time-window for each meerkat. After plotting histograms of log VeDBA (always bimodal), we manually determined the appropriate threshold value to demarcate low- and high-activity levels. New behavioral sequences based on VeDBA-thresholding were obtained in this way. The three main analyses from the main text: bout duration distributions, hazard functions, and predictivity decay curves were all computed for these new behavioral sequences.

We found that all our results from the main text were successfully replicated in the alternate behavioral sequences (Figure 6): bout duration distributions were consistently heavy-tailed (with truncated power-law best fits in all instances and behavioral states), hazard functions of bouts of both high- and low-activity states were decreasing, and predictivity decay continues to fit best to a truncated power-law.

This set of additional analyses demonstrates the robustness of our results to the choice of time-window length as well as to any artifacts caused by our classifiers. In the main text, meerkat results

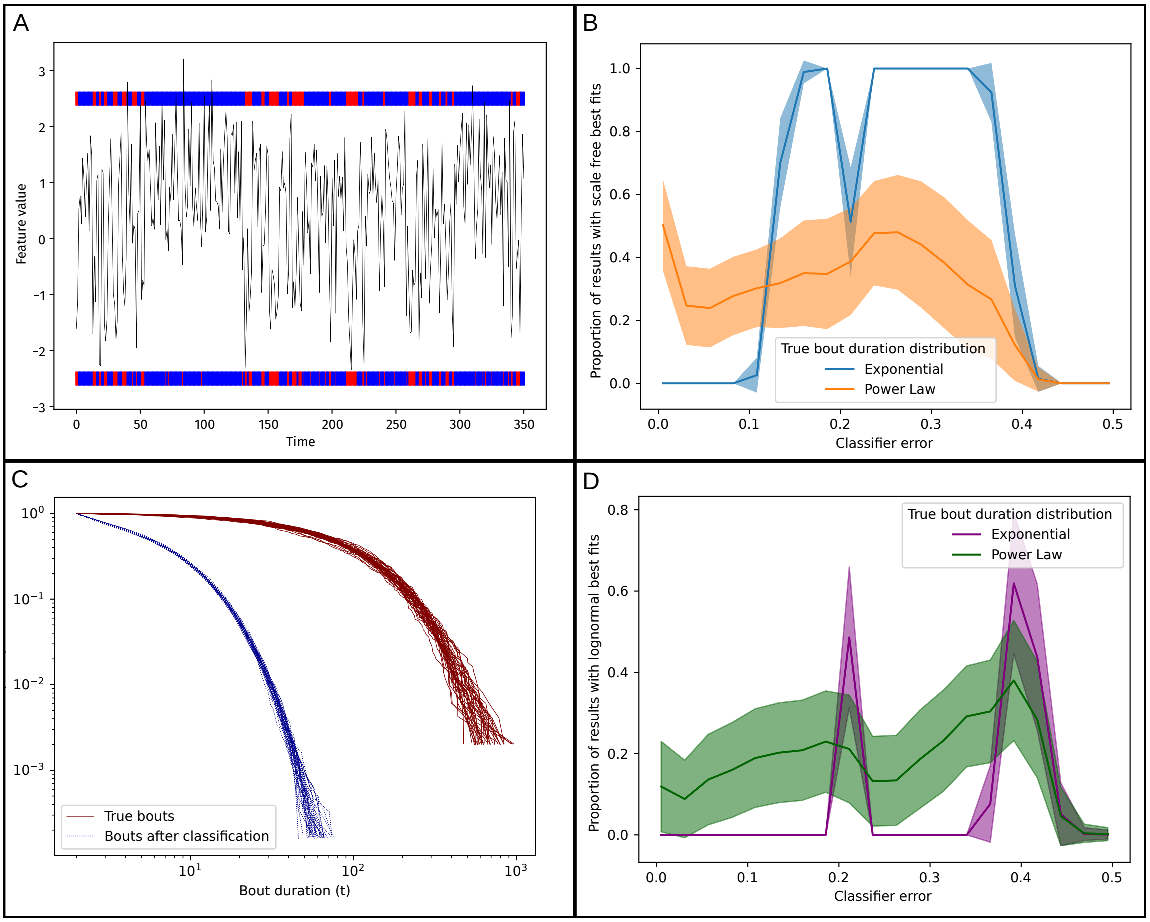

**Figure 3:** (A) Illustration of a simulation: true behavioral sequences (band on top) were used to generate a time-varying feature (black line), which was then used to compute the perceived behavioral sequence (bottom band). (B) Fitting distributions to the perceived bouts, we saw that power-laws and truncated power-laws are destroyed by error-prone classifiers as expected. However, at intermediate error rates, truncated power-laws were spuriously almost always detected in memory-less sequences. (C) At an error rate of 0.25, long bouts ($> 100$ s) are destroyed almost completely. Shorter bouts are affected lesser by errors. This phenomenon could lead to exponentially distributed data having a truncated power-law best fit, due to which a heavy tail is detected in a memory-less process. Nonetheless the perceived distribution's tail is not as heavy as that of the true distribution. (D) At all error values, power-law bout duration distributions could be perceived as lognormal. However, exponential bout duration distributions are only perceived as lognormal at specific error rates.

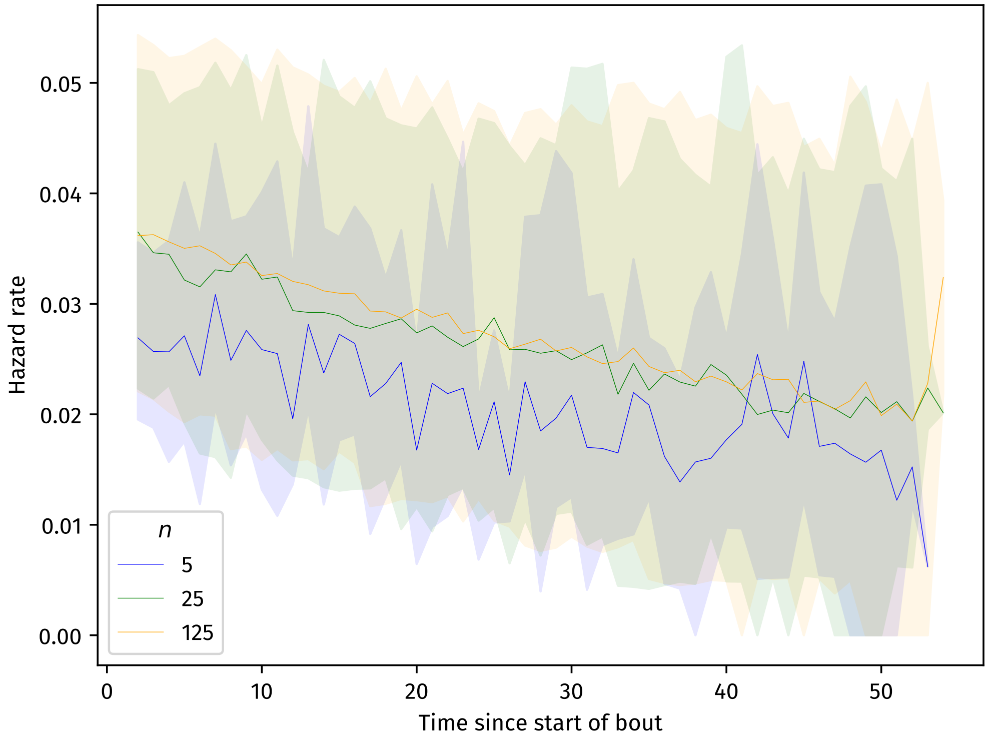

**Figure 4:** Average hazard functions of behavioral bouts in groups of 5 (blue), 25 (green) and 125 (orange) identical socially reinforcing individuals. The hazard function decreases in all cases, seemingly showing some kind of positive feedback. This apparent positive feedback comes from the contextual dependence of bout lengths on the proportion of group members performing the same behavior.

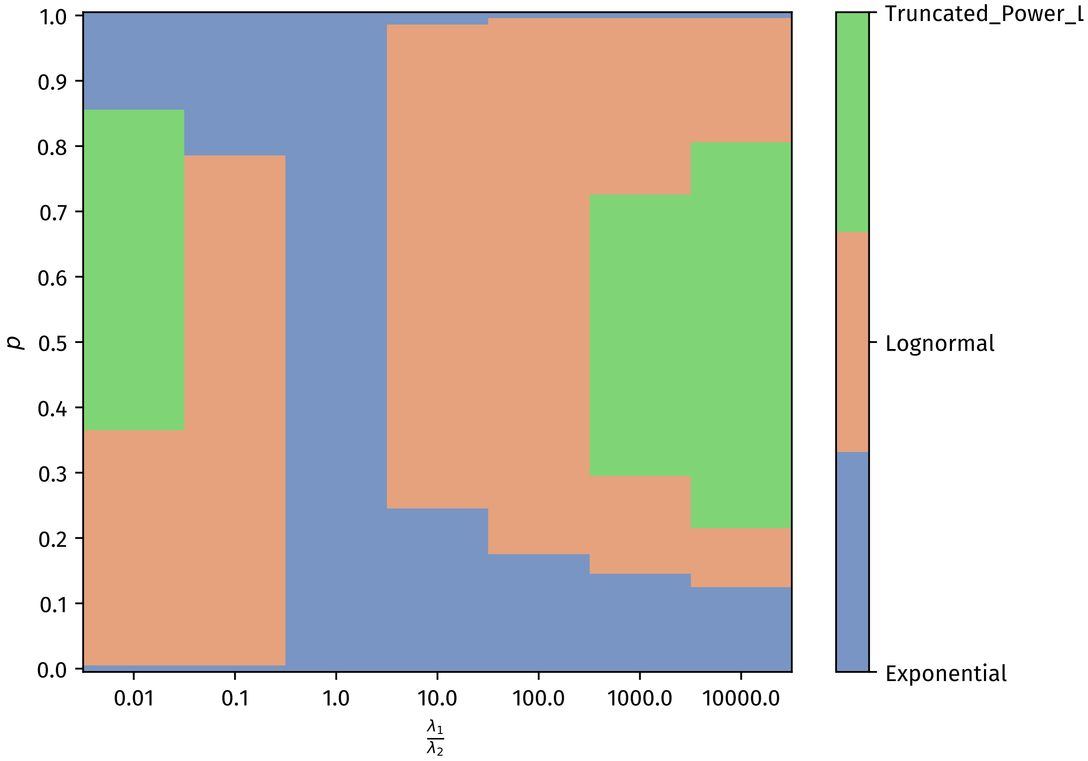

**Figure 5:** Most common best fit distributions out of 100 for each combination of $p$, $\lambda_1$, and $\lambda_2$. We see that specific regions of the parameter space have specific best-fit distributions. When both distributions are identical ($\lambda_1 = \lambda_2$), an exponential best fit is observed as expected. Specific areas of the parameter space with similar proportions of dissimilar exponentials (i.e., when the tail of one distribution decays much faster than the tail of another) fit best to truncated power-law distributions. Lognormal best-fits were most common across parameter space. Power-laws without truncation did not fit data generated from any pair of exponentials.

**Table 2:** Best fit distributions for each state for each individual (on each row), for (a) meerkats, (b) coatis, and (c) hyenas. Distribution fits are abbreviated (TPL: Truncated Power-law and LN: Lognormal) Distribution fits were determined by AICs (see Materials and Methods in the main text).

(a)

| Running | Foraging | Vigilance |
|---|---|---|
| Insuff. data for fitting | TPL($\alpha$=1.356; $\lambda$=3.6 × $10^{-4}$) | LN($\mu$=0.181; $\sigma$=1.606) |
| | TPL($\alpha$=1.339; $\lambda$=2.3 × $10^{-4}$) | LN($\mu$=0.616; $\sigma$=1.226) |
| | TPL($\alpha$=1.475; $\lambda$=1.6 × $10^{-4}$) | LN($\mu$=-0.221; $\sigma$=1.780) |
| | LN($\mu$=0.721; $\sigma$=2.605) | LN($\mu$=0.918; $\sigma$=1.811) |
| | TPL($\alpha$=1.212; $\lambda$=5.5 × $10^{-4}$) | LN($\mu$=0.612; $\sigma$=1.636) |
| | LN($\mu$=0.732; $\sigma$=2.533) | LN($\mu$=1.230; $\sigma$=1.285) |
| | TPL($\alpha$=1.239; $\lambda$=4.0 × $10^{-4}$) | LN($\mu$=0.200; $\sigma$=1.884) |
| | LN($\mu$=1.694; $\sigma$=2.433) | TPL($\alpha$=1.389; $\lambda$=0.010) |
| | TPL($\alpha$=1.340; $\lambda$=2.5 × $10^{-4}$) | LN($\mu$=0.797; $\sigma$=1.558) |
| | TPL($\alpha$=1.465; $\lambda$=1.5 × $10^{-4}$) | TPL($\alpha$=1.540; $\lambda$=0.067) |
| | LN($\mu$=2.132; $\sigma$=2.672) | TPL($\alpha$=1.456; $\lambda$=0.006) |
| | TPL($\alpha$=1.508; $\lambda$=1.0 × $10^{-4}$) | LN($\mu$=0.267; $\sigma$=1.149) |
| | LN($\mu$=-0.468; $\sigma$=2.975) | LN($\mu$=0.840; $\sigma$=1.455) |
| | TPL($\alpha$=1.242; $\lambda$=6.1 × $10^{-4}$) | LN($\mu$=0.551; $\sigma$=1.671) |
| | TPL($\alpha$=1.380; $\lambda$=1.8 × $10^{-4}$) | TPL($\alpha$=1.076; $\lambda$=0.058) |

(b)

| Low | High |
|---|---|
| LN($\mu$=-3.480; $\sigma$=2.707) | TPL($\alpha$=1.321; $\lambda$=0.003) |
| LN($\mu$=-4.398; $\sigma$=2.631) | TPL($\alpha$=1.234; $\lambda$=0.003) |
| LN($\mu$=-0.620; $\sigma$=1.907) | TPL($\alpha$=1.429; $\lambda$=0.003) |
| LN($\mu$=-1.694; $\sigma$=2.059) | LN($\mu$=0.184; $\sigma$=2.370) |
| LN($\mu$=-1.032; $\sigma$=2.041) | TPL($\alpha$=1.400; $\lambda$=0.002) |
| LN($\mu$=-0.540; $\sigma$=1.844) | TPL($\alpha$=1.288; $\lambda$=0.003) |
| LN($\mu$=-2.334; $\sigma$=2.691) | TPL($\alpha$=1.302; $\lambda$=0.003) |
| LN($\mu$=-3.195; $\sigma$=2.353) | TPL($\alpha$=1.272; $\lambda$=0.003) |
| LN($\mu$=-2.723; $\sigma$=2.392) | TPL($\alpha$=1.272; $\lambda$=0.003) |

(c)

| LYUP | WALK | LOPE | STAND | LYING |
|---|---|---|---|---|
| LN($\mu$=-1.770; $\sigma$=1.667) | TPL($\alpha$=1.496; $\lambda$=0.017) | TPL($\alpha$=1.000; $\lambda$=0.047) | LN($\mu$=-0.365; $\sigma$=1.384) | TPL($\alpha$=1.258; $\lambda$=0.002) |
| LN($\mu$=-3.299; $\sigma$=1.946) | TPL($\alpha$=1.264; $\lambda$=0.022) | LN($\mu$=1.219; $\sigma$=1.087) | LN($\mu$=-0.379; $\sigma$=1.385) | TPL($\alpha$=1.057; $\lambda$=0.004) |
| TPL($\alpha$=2.325; $\lambda$=1 × $10^{-4}$) | TPL($\alpha$=1.560; $\lambda$=0.014) | LN($\mu$=1.374; $\sigma$=1.052) | LN($\mu$=0.354; $\sigma$=1.272) | TPL($\alpha$=1.266; $\lambda$=0.003) |
| LN($\mu$=-3.792; $\sigma$=2.490) | TPL($\alpha$=1.566; $\lambda$=0.018) | LN($\mu$=1.224; $\sigma$=1.177) | LN($\mu$=-0.053; $\sigma$=1.349) | TPL($\alpha$=1.355; $\lambda$=0.002) |
| LN($\mu$=-5.252; $\sigma$=2.271) | TPL($\alpha$=1.417; $\lambda$=0.019) | TPL($\alpha$=1.060; $\lambda$=0.074) | LN($\mu$=-0.382; $\sigma$=1.440) | TPL($\alpha$=1.282; $\lambda$=0.004) |

**Table 3:** Parameter and $R^2$ values computed for predictivity decay in (a) meerkats, (b) coatis, and (c) hyenas. Each row represents one individual. In all cases, Adjusted Mutual Information was found to decay as an exponentially truncated power-law.

| | $m$ | $\alpha$ | $\lambda$ | $R^2$ **exponential** | $R^2$ **powerlaw** | $R^2$ **truncated powerlaw** |
|---|---|---|---|---|---|---|
| (a) | 0.566 | 0.226 | 0.0016 | 0.902 | 0.9504 | 0.9987 |
| | 0.616 | 0.185 | 0.0015 | 0.9270 | 0.9075 | 0.9991 |
| | 0.563 | 0.046 | 0.0075 | 0.8910 | 0.7396 | 0.8934 |
| | 0.699 | 0.253 | 0.0009 | 0.8491 | 0.9626 | 0.9972 |
| | 0.765 | 0.169 | 0.0032 | 0.9515 | 0.9170 | 0.9979 |
| | 0.698 | 0.216 | 0.0021 | 0.9188 | 0.9506 | 0.9916 |
| | 0.471 | 0.219 | 0.0008 | 0.8845 | 0.9541 | 0.9774 |
| | 0.647 | 0.293 | 0.0021 | 0.8871 | 0.9686 | 0.9972 |
| | 0.669 | 0.156 | 0.0025 | 0.9499 | 0.9052 | 0.9991 |
| | 0.711 | 0.225 | 0.0017 | 0.9090 | 0.9506 | 0.9977 |
| | 0.717 | 0.164 | 0.0027 | 0.9496 | 0.9122 | 0.9983 |
| | 0.716 | 0.200 | 0.0056 | 0.9578 | 0.9298 | 0.9956 |
| | 0.689 | 0.213 | 0.0011 | 0.8866 | 0.9537 | 0.9965 |
| | 0.763 | 0.181 | 0.0031 | 0.9477 | 0.9210 | 0.9973 |
| | 0.810 | 0.205 | 0.0009 | 0.8876 | 0.9501 | 0.9944 |
| | $m$ | $\alpha$ | $\lambda$ | $R^2$ **exponential** | $R^2$ **powerlaw** | $R^2$ **truncated powerlaw** |
| (b) | 0.5434 | 0.1762 | 0.0005 | 0.8487 | 0.9403 | 0.9975 |
| | 0.5621 | 0.1644 | 0.0005 | 0.8439 | 0.9382 | 0.9972 |
| | 0.5347 | 0.1594 | 0.0005 | 0.8564 | 0.9356 | 0.9966 |
| | 0.6689 | 0.1263 | 0.0008 | 0.9171 | 0.8963 | 0.9960 |
| | 0.5983 | 0.1424 | 0.0003 | 0.7899 | 0.9469 | 0.9962 |
| | 0.5986 | 0.1599 | 0.0004 | 0.8354 | 0.9457 | 0.9959 |
| | 0.5415 | 0.1650 | 0.0005 | 0.8472 | 0.9433 | 0.9940 |
| | 0.5802 | 0.1720 | 0.0003 | 0.8142 | 0.9560 | 0.9976 |
| | 0.5626 | 0.1592 | 0.0003 | 0.7960 | 0.9439 | 0.9932 |
| | $m$ | $\alpha$ | $\lambda$ | $R^2$ **exponential** | $R^2$ **powerlaw** | $R^2$ **truncated powerlaw** |
| (c) | 0.6241 | 0.1921 | 0.0008 | 0.8760 | 0.9462 | 0.9988 |
| | 0.6488 | 0.1673 | 0.0015 | 0.9257 | 0.9182 | 0.9992 |
| | 0.5874 | 0.1847 | 0.0013 | 0.9089 | 0.9318 | 0.9985 |
| | 0.6347 | 0.1874 | 0.0014 | 0.9141 | 0.9328 | 0.9985 |
| | 0.6474 | 0.2057 | 0.0014 | 0.9079 | 0.9432 | 0.9986 |

**Table 4:** $\alpha_{DFA}$ values for each behavior for (a) meerkats, (b) coatis, and (c) hyenas. Each row shows data for one individual.

(a)

| Vigilance or resting | Foraging | Running |
|---|---|---|
| 1.0925 | 1.0914 | 0.9698 |
| 1.0985 | 1.0984 | 0.9516 |
| 1.1354 | 1.1348 | 0.9586 |
| 1.1406 | 1.1401 | 0.9344 |
| 1.1375 | 1.1369 | 0.9624 |
| 1.1710 | 1.1704 | 0.9446 |
| 1.1373 | 1.1366 | 0.9542 |
| 1.1738 | 1.1728 | 0.9248 |
| 1.1306 | 1.1301 | 0.9701 |
| 1.0813 | 1.0809 | 0.9556 |
| 1.1058 | 1.1056 | 0.9606 |
| 1.1414 | 1.1399 | 0.9455 |
| 1.0701 | 1.0696 | 0.9350 |
| 1.0671 | 1.0666 | 0.9627 |
| 1.1216 | 1.1211 | 0.9456 |

(b)

| Low activity | High activity |
|---|---|
| 1.1145 | 1.1145 |
| 1.0889 | 1.0889 |
| 1.1040 | 1.1040 |
| 1.1169 | 1.1169 |
| 1.1027 | 1.1027 |
| 1.0978 | 1.0978 |
| 1.1095 | 1.1095 |
| 1.1950 | 1.1950 |
| 1.1058 | 1.1058 |

(c)

| Lying (head down) | Walking | Standing | Lying (head up) | Running |
|---|---|---|---|---|
| 1.1708 | 1.0950 | 0.9731 | 0.9349 | 1.0554 |
| 1.1465 | 1.0505 | 0.9837 | 1.0122 | 1.0631 |
| 1.1967 | 1.1055 | 1.0069 | 0.9613 | 1.0689 |
| 1.1453 | 1.0609 | 0.9903 | 0.9738 | 1.0438 |
| 1.1610 | 1.0513 | 0.9717 | 0.9389 | 1.0510 |

are based on behavioral sequences inferred with 1 s time-windows using a supervised machine-learning approach (section 1). 1 s and 2 s time-windows are both effective in distinguishing separate meerkat behaviors from accelerometer data, and the behavioral sequences obtained in both cases thus retain the true dynamics of behavior.

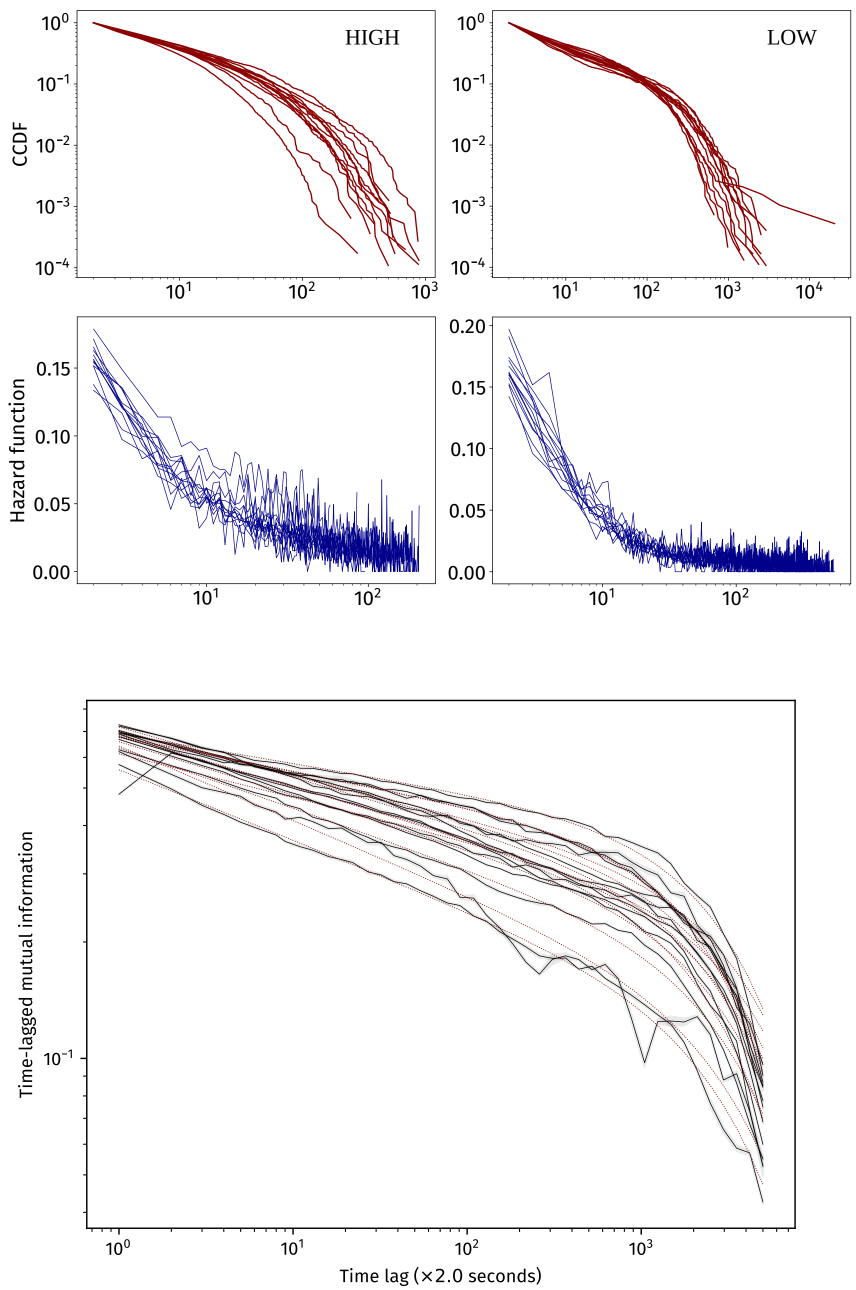

**Figure 6:** Repeat analysis of meerkat behavioral dynamics with alternate, unsupervised learning-based approach for behavior inference. Behavioral sequences are inferred from accelerometer data with 2 s time-windows. All results from the main-text are replicated. (i) bout duration distributions are consistent and heavy-tailed for all individuals (row 1, red) for both activity levels; (ii) hazard functions are consistently decreasing for all individuals (row 2, blue) in both activity levels; and (iii) mutual information decay follows a truncated power-law across individuals.
